# Description of the structural and functional genomic composition of *L. santarosai* serovar Alice

**DOI:** 10.1101/2024.08.10.607426

**Authors:** Rafael Guillermo Villarreal-Julio, Valeria Isabel Manjarrez Montes, Brayan Estiven Ordoñez Perez, Jonny Andrés Yepes-Blandón

## Abstract

In Colombia, pathogenic knowledge of endemic *Leptospira* species is limited. *Leptospira* santarosai serogroup Autumnalis serovar Alice Strain JET was isolated from a patient from Urabá Antioquia, so it is necessary to know more about its genomic components and pathogenomics. A comparative analysis of the bacterial genome was carried out by computational biology. Complete information for analysis was obtained fromhttps://www.ncbi.nlm.nih.gov/from where the complete genome sequences of 67 species of the genus *Leptospira* spp were accessed. The genome of the Colombian strain consists of a major chromosome (chromosome I) and the minor chromosome (chromosome II), which add up to a total of 4.13 Mb. The chromosomes have a G+C percentage of 41.6% three genes for ribosomal RNA distributed 1 for the 23s subunit, 1 for the 16s subunit and 1 for the 5s subunit, 6 for other types of RNA and 37 transfer RNA genes. Two chromosomal pseudomolecules were inferred for the Colombian strain, assigning proteins 3,796 and 318 to chromosomes I and II respectively. Genomic identity analysis resulted in a nucleotide identity of 92.95% and a high degree of synteny between chromosomes. 53 and 68 exclusive proteins were identified for serovar Shermani and the Colombian strain respectively. Of the 68 proteins, most are hypothetical, however, nine transposases, two integrases and one membrane protein were identified. The Colombian strain shares orthologous proteins with members of the different subgroups of species that range between 3,286-2,600 with pathogenic species, 2,416-2,296 intermediate species and 1,972-1,864 saprophytic species and 40.98% of the proteins present in the genome of The Colombian strain has a high percentage of identity with the proteins of the main pathogenic species *L. interrogans* (80-100% similarity), which demonstrates a strong genetic relationship between the Colombian strain and the pathogenic subgroup. For the Colombian strain, it was found to contain CRISPR/Cas repeats and 9 families of class 1, type I and subtype IA and IE were located where each of these subtypes contained 8 cas genes; In addition, 25 proteins were found involved in the de novo biosynthesis of vitamin B12. In the Colombian strain, three species of different phage structures were found that could facilitate infection and subsequent colonization in the host. Therefore, the objective is to describe the structural and functional genomic composition of *L. santarosai* serovar Alice. The genomic characteristics evidenced are basic in the knowledge of the bacterial mechanisms, the evolutionary, phylogenetic and pathogenic analysis of *Leptospira* spp that affects the population of Colombian endemic areas.

## Introduction

leptospirosisIt is the most frequently reported zoonosis worldwide,neglected reemergence, which is classified as a potentially fatal febrile syndrome in humans who are accidental hosts. In some patients with severe cases, the disease produces acute renal failure, liver injury or pulmonary hemorrhage syndrome (1) and is caused by gram-negative spirochetes belonging to the genus *Leptospira*, which has a wide genetic diversity (1–4). The genus comprises 67 species that are subdivided into groups of pathogenic, intermediately pathogenic, and free-living saprophytic species, which are non-pathogenic and live in the environment (5). The three groups present profound diversity in terms of their distribution in the environment, transmission mechanisms and genetic composition (4).

This diversity creates a challenge for the search for genomic targets and for the knowledge of virulence factors of *Leptospira* spp. For those who have been interested in microbiology, one of the great contributions that has been obtained from the emerging analysis of genomes is to understand the biological mechanisms by which a microbial genus, such as *Leptospira*, survives in a wide profile of microenvironments, infects a wide variety of hosts, how it is phylogenetically structured and how it can become virulent. This knowledge enriches the biological information of the genus through the contributions of comparative genomics.

Although the genus has been recognized for more than a century worldwide (6), only in the late 1990s were genetic, genomic and molecular mechanisms established to understand this group of spirochetes from the perspective offered by the known emerging science. as pathogenomics (7). The investigation of the pathogenesis mechanisms of *Leptospira* has been difficult due, among other reasons, to the difficulty of maintaining viable cultures of pathogenic leptospires given their condition as a “fastidius” bacterial group, with slow growth and poor transformation capacity with genetic tools to manipulate genome.

Fortunately, as a result of new genomic sequencing methodologies in the decade from 2010 to 2020, genomes of *Leptospira* spp strains are available, which have allowed an increase in knowledge of the pathogenesis of this bacterial genus through the analysis of functional genomics. and as a result of the above, virulence factors have been inferred (8). The contribution of this knowledge generation strategy allowed the understanding of many pathogenic aspects, such as evidencing the dynamic and diverse nature of the microbial group, and in 2019, 67 species were recognized (5.9), when in the nineties only L. biflexa and *L. interrogans* were included in the genus.

In Colombia, knowledge through pathogenomics of endemic *Leptospira* species that affect human health is limited (10–13). Thanks to a collaboration of our group, *L. santarosai* serogroup Autumnalis serovar Alice Strain JET was the first Colombian strain included in 2012 in the J. Craig Venter Institute project (The global *Leptospira* Genome Project - US National Institute of Allergy and Infectious Diseaseshttps://www.jcvi.org/research/Leptospira-genomics-and-human-health). Since that date, its genome has been available, along with the 1,257 other genomes registered in the genus to date, at the National Center of Biotechnology Information –NCBI.

*L. santarosai*serogroup Autumnalis serovar Alice Cepa JET was isolated in 2008 from a patient with undifferentiated febrile syndrome from one of the regions where the most cases of leptospirosis are reported in Colombia, Urabá Antioquia (10). After its taxonomic determination, it has been studied in some biological and genomic aspects (10,12); however, it is necessary to know more about its genomic composition and the mechanisms of its pathogenomics, based on a description of its similarities and particularities compared to other serovars of both the genus and the species L. santarosai. To respond to this need, the study was proposed seeking to ensure that the knowledge generated has application in the functional genomics of the Colombian strain of *Leptospira* using microbial pathogenomics. The contribution to knowledge was defined to answer the question: What is the genomic structure of a Colombian strain of *L. santarosai* determined by a comparative genomic analysis of pathogenic, intermediate and saprophytic species of the genus *Leptospira*?

Knowing and analyzing the genome sequence of pathogenic *Leptospira* species facilitates the localization of their genes and the study of the function of several genes necessary in tissue degradation, invasion, dissemination, evasion of the immune response and adaptation to the environment and to the host, as well as the characterization of sequences that regulate the genetic expression of its antigenic proteins (14). The growing increase in information generated from genome sequencing projects has led to the advancement of new disciplines such as computational biology and bioinformatics, which use computational tools to organize, store, examine and compare sequences with the purpose of identifying similarities to others. biological systems to thus define functions through pathogenic analysis (4). With bioinformatics tools, the study of genes that are common among pathogenic strains makes it possible to perform comparative genomics that allows identifying genetic factors necessary in the bacteria to infect and cause the disease (15,16). Thus, performing the comparative and functional genetic analysis of *L. santarosai* serogroup Autumnalis serovar Alice through bioinformatics models is important because it specifically provides a better understanding of the genetic bases of the virulence and adaptation mechanisms to both the environment and the environment. host.

## Methodology

### Methodological approach and type of study

The research is a quantitative approach of a descriptive type of analysis of the bacterial genome, where numerical measurement of data obtained through computational biology was carried out. Complete information for bioinformatic analyzes was obtained from the database stored at the National Center for Biotechnology Information (NCBI) [Internet]. Bethesda (MD): National Library of Medicine (US), National Center for Biotechnology Information; [1988] – [cited 2018 Dec. 10]. Available from:https://www.ncbi.nlm.nih.gov/from where the sequences of complete genomes, nucleotides, total genes of interest, proteins and proteome of bacteria of the genus *Leptospira* spp.

The genomic data of the 67 species that make up the three groups of pathogenic, intermediate and saprophytic *Leptospira* reported up to that moment and for which genomic information was available in the databases in GenBank were included. The Colombian strain under study was *Leptospira* santarosai serogroup Autumnalis serovar Alice Cepa JET (GenBank assembly accession GCA_000306475.2 -NCBI: txid1193057).

For some analyzes related to orthology between the 3 groups, 4 reference species that had a complete genome were included. The genomic comparison was carried out with them; these were *Leptospira* santarosai serovar Shermani strain LT 821 (GenBank assembly accession GCA_000313175.2), *Leptospira* interrogans serovar Copenhageni strain Fiocruz L1-130 (GenBank assembly accession GCA_000007685.1), *Leptospira* licerasiae serovar Varillal strain VAR 010 (GenBank assembly accession GCA_0002 44755.3) and *Leptospira* biflexa serovar Patoc strain ’Patoc 1 (Paris)’ (GenBank assembly accession GCA_000017685.1), these in turn represent the three subgroups, pathogenic, intermediate and saprophytic and are included within the group of the 67 species of *Leptospira* mentioned. .

#### Colombian strain of *Leptospira*

The strain *L. santarosai* serogroup Autumnalis serovar Alice of Colombian origin was isolated in 2008 from a male patient from Apartadó, Antioquia, Colombia, who had been suffering from fever for three days and who had a blood culture performed in the middle Fletcher’s semisolid. This culture was replicated and maintained as a laboratory isolate. Said isolate was typed using the 16s ribosomal gene and MLST, for the identification of the species and serovar, considering it as a laboratory reference strain (3). Since then, the bacterial subcultures were maintained in EMJH liquid medium (Ellinghausen-McCullough-Johnson-Harris, BectonDickinson-Biosciences) supplemented with 10% commercial enrichment medium (Becton-Dickinson-Biosciences), incubated between 26°C and 30°C. under aerobic conditions and frozen at -80°C following standard methodologies (17).

#### Genome sequencing of the Colombian strain, assembly and annotation

Thanks to a collaboration of our group, *L. santarosai* serogroup Autumnalis serovar Alice Strain JET was the first Colombian strain included in 2012 in the J. Craig Venter Institute project (The global *Leptospira* Genome Project - US National Institute of Allergy and Infectious Diseaseshttps://www.jcvi.org/research/Leptospira-genomics-and-human-health<u>). The</u>Sequencing methodology used was: dual next-generation sequencing (NGS) platforms 454 GS FLX Titanium (Roche, Branford, USA) and Illumina HiSeq 2000 (Illumina Inc., San Diego, CA, USA). UU), to obtain better coverage of the genome sequence. An RS Pacific Biosciences sequencer (PacBio; Pacific Biosciences, Menlo Park, USA) was used.

De novo assembly was performed using Celera Assembler v. 7.0. The order and orientation of these contigs were corroborated using an optical mapping system (Opgen Technologies Inc., Madison, WI, USA).

DNA and/or bacterial genomic material are available in the BEI resource repository (http://beiresources.org). This sequencing project was funded with federal funds from the National Institute of Allergy and Infectious Diseases of the National Institutes of Health, Department of Health and Human Services through the Genomic Sequencing Centers for Infectious Diseases under contract number HHSN272200900007C. The sequence was entered into the NCBI platform by the J. Craig Venter Institute and the annotation of the contigs was updated as of March 2013. The genome sequence of the Colombian strain is found in the GenBank database of the NCBI platform with the accession number NZ_AKWS00000000.

#### Downloading genomes, CDS and proteomes

The genomes of the 67 *Leptospira* species, CDS and their respective annotated proteins were downloaded from the NCBI database. These species represent pathogenic, intermediate and saprophytic subgroups.

#### Analysis of genomic identity between the Colombian strain and the reference genome of L. santarosai

Genomic identity analysis was performed using the genome of the Colombian strain and the reference genome of *Leptospira* santarosai serovar Shermani. Assignment of contigs to reference chromosomes was performed using the NUCmer bioinformatics tool of the MUMmer v.3.1 package. Subsequently, the pseudo-molecules of chromosomes I and II were inferred using the show-tiling option, using the same package. The synthesis analysis between reference chromosomes and pseudochromosomes was visualized by the bioinformatics tools of Mauve and BRIG (11,12,34).

#### Confirmation and update of the genomic and characterization of L. santarosai serovar Alice used phylogenetic analysis of the 16s ribosomal RNA gene

For molecular phylogenetic confirmation and update with the previous and new described species of the genus *Leptospira* spp, the sequence of the 16s ribosomal gene belonging to the 67 species that make up the genus *Leptospira* was used. The sequences were acquired from the NCBI database except for *L. interrogans* and L. alstonii, which were extracted using the Artemis viewer from their respective annotated genome. These sequences were aligned through the ClustalW program. Subsequently, this alignment was used to make phylogenetic trees using the MEGAX program, where the Neighbor-Joining method was used with 1000 bootstrap replicates and calculating the evolutionary distances by the Kimura-2 parametric method. The NCBI accession ID of the 16s ribosomal gene sequences used for this analysis can be displayed in the phylogenetic tree image (Fig. 1).

**Figure 1.**
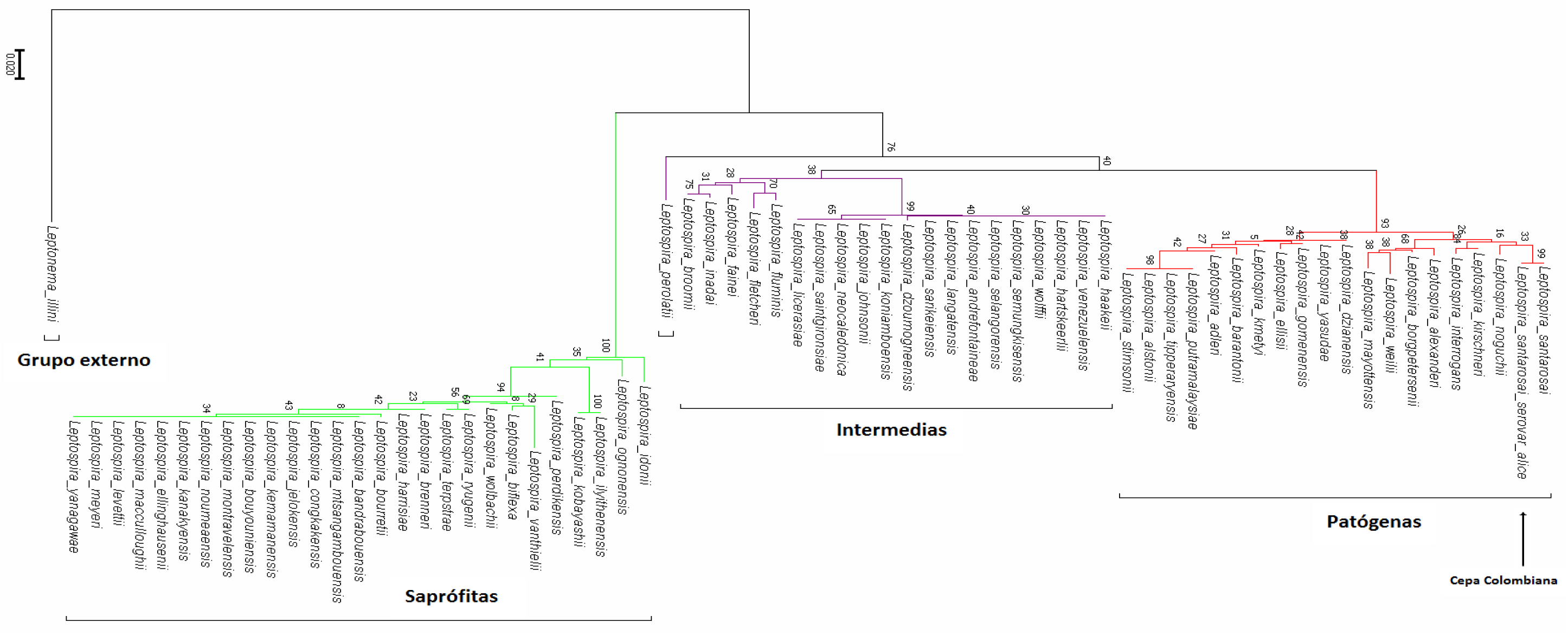
Phylogenetic characterization of the Colombian strain against the described *Leptospira* species: The strain was located with complete similarity to *L. santarosai* as indicated by the arrow. The phylogenetic analysis of the 16s ribosomal gene is presented through the MEGA7 program through the neighbor-joining algorithm using the Bootstrap model of 1000 repetitions and the 2-parameter kimura parameter. The figure shows for the first time worldwide the separation of the 67 species of *Leptospira* hats now reported into groups of pathogenic (P1), intermediate pathogenic (P2) and saprophytic (S1 and S2) species marked by red, purple and green colors respectively. . In black you can see the external group that includes Leptonema, a bacteria different from the genus *Leptospira*.The figure shows the inability of the 16s ribosomal gene to differentiate, for example, between L. putramalaysiae/L. tipperaryensis/L. alstoni/L. stimsonii,. terpstrae/L. ryugenii, L. meyeri/L. macculloughii/L. levettii/L. yanagawae, L. licerasiae/L. saintgironsiae / L. neocaledonic, and L. venezuelensis / L. haakeii / L. hartskeerlii / L. wolffii among others. Own elaboration by the author of this work (48).

#### Phylogenetic distribution of isolates belonging to L. santarosai

To identify serovars related to L. santarosai, two genes were chosen with the following characteristics: conserved in the genus, highly polymorphic and with the largest number of sequences reported in the NCBI database. Thus, the gyrB and secY genes were chosen to differentiate serovars related to L. santarosai, and these genes were used in a concatenated sequence analysis to identify 34 serovars.

The haplotype network of the *L. santarosai* isolates with the gyrB and secY genes was carried out using NCBI Tree Viewer (Treeviewer CGI version: 1.17.1.r40315) using 30 sequences of *L. santarosai* isolates, which made up all the isolates described worldwide. The Tree Viewer program was implemented using the ExtJS JavaScript library.

#### Genome characteristics of the Colombian strain L. santarosai serovar Alice and the 67 species of Leptospira

The characteristics of all reference species were downloaded from the genomes available from Genbank at NCBI and tabulated in a table generated in Excel where the main structural characteristics were described such as *Leptospira* species name, size (Mb), GC% (percentage of guanine and cytokine), number of proteins, rRNA (ribosomal RNA), tRNA (transfer RNA), Other RNAs, number of genes, number of pseudogenes and finally the NCBI accession number of each of them.

#### Concatenated sequence analysis to identify serovars related to L. santarosai serovar Alice

Concatenated sequence analysis was performed using the gyrB and secY genes, which are constitutively expressed and highly polymorphic among serovars related to *L. santarosai* (18,19). These genes were downloaded from the NCBI database and aligned using the ClustalX program (18). To investigate the phylogenetic relationship between serovars related to L. santarosai, a median linkage network analysis was done. The network was performed using popart 4.6.1.2 (19) using concatenated sequences of secY and gyrB genes (18,19).

#### Comparative genomic analysis to determine the core genome and pangenome of the Colombian strain among different Leptospira species

Comparative genomic analysis was performed using the genome of the Colombian strain, comparing it with the 67 genomes of the genus *Leptospira* previously downloaded from NCBI. These species represent pathogenic, intermediate and saprophytic subgroups (Fig. 6). The OrthoMCL algorithm was used to cluster proteins into orthologous groups and unique genes among species from the different groups of pathogenic, intermediate, and saprophytic species based on their sequence identity (20), and the InParanoid algorithm was used to identify paired orthologous proteins (21).

The sequences in FASTA format of the three groups created from their association with pathogenicity were filtered following protocols established by the OrthoMCL program, in which a Markov clustering algorithm was used to group orthologs and paralogs (putative) (20). The genes were identified and mapped through the GI code of the NCBI database. The results were grouped based on the presence of unique genes per pathogenicity category or by the presence of genes exclusively in certain categories and absence in the rest. The analysis was used to identify genes associated with the pathogenicity of bacteria and to classify genes and identify groups of orthologs in several categories: group 1. Genes that reported at least one ortholog with another species, group 2. Genes from pathogens that had no orthologs in non-pathogenic groups, group 3. Genes that did not have orthologs in non-pathogenic groups and reported orthologs in all pathogenic and intermediate species and group 4. Total unique genes among all species.

For group 3 of pathogen genes that did not have orthologs in non-pathogenic groups, that is, genes that were found only in pathogens, the cellular location of each of them was also sought; For this purpose the proteins were mapped to the UniProt database (www.uniprot.org) and each of the genes found were searched for annotated information from the GeneOntology in its sub-cellular location component. The search for orthologs and unique genes between pathogenic and intermediate species was also carried out using the same methodology.

#### Graphical description of the core genome through the distribution of orthologous proteins among 67 species of the genus Leptospira

Common or unique orthologous proteins between the genomes were plotted via the Orthovenn page (http://www.bioinfogenome.net/OrthoVenn/). For the graph, representative species of each group were used, thus for the pathogenic group the pathogenic species *L. santarosai* serovar Shermani strain LT 821 were used to compare it with the Colombian strain *L. santarosai* serovar Alice in addition to *L. interrogans* serovar Copenhageni strain Fiocruz L1 -130, for the intermediate group *L. licerasiae* serovar Varillal strain VAR 010, for the saprophytic group *L. biflexa* serovar Patoc strain ’Patoc 1 (Paris)’.

A similar analysis was performed to detect orthologous proteins between *L. santarosai* serovar Alice and all 29 strains of the *L. santarosai* species and for this reason the common or unique orthologous proteins between the genomes were plotted through the Orthovenn page (http://www.bioinfogenome.net/OrthoVenn/).

Finally, all proteins reported for each species and serovar in the NCBI database were downloaded in fasta format. A global protein similarity search between genomes was performed using ggsearch36 v.36.3.6 software, as implemented in the Fasta36 package (21).

#### Synteny analysis carried out with the MAUVE program between serovars Shermani and Alice belonging to the species L. santarosai

Synteny analysis between reference chromosomes and pseudochromosomes of *L. santarosai* serovar Alice and representative species of the three subgroups of the *Leptospira* genus was performed with the ACT application. The creation of the comparison file was done by terminal in Biolinux.

#### Analysis of the pangenome and core genome of the Colombian strain L. santarosai and different pathogenic, intermediate and saprophytic Leptospira species

Taking into account the *Leptospira* species that are currently sequenced in complete form or in contigs, a search of the pangenome and core genome of the studied species was carried out. For this, the PanOCT program was implemented (http://sourceforge.net/p/panoct/home/Home/), which is a Perl script that not only looks for strategies to detect orthologs by conventional methods, but also takes into account the specific position in the genome of the coding sequences and the conservation of neighboring genes.

To work satisfactorily with the program, it was necessary to create different files, taking into account protein information, such as the annotation files of the coding sequences (CDS) according to the Genbank format initially downloaded for each of the strains. The specific files that had to be created are the following: Multiple alignment between all strains through the Blastp package of the NCBI BLAST+ toolkit (http://blast.ncbi.nlm.nih.gov/Blast.cgi), identifiers of each of the strains subjected to the algorithm, attribute file in which the contig is specified, the position in the genome, the ID and the function of the protein belonging to each of the strains analyzed and finally the protein sequences in FASTA format.

#### *Unique proteins found in the Colombian species L. santarosai* serovar Alice

Annotations were performed using Rapid Annotations using Subsystems Technology (RAST) version 4.0. The genome was uploaded to GenBank® format as a set of contigs in FASTA format, complete annotation typically occurred within 12 to 24 hours, subsequently the annotated genome was compared to the hundreds of genomes maintained within the SEED integration in a internal base. Subsequently, the unique proteins option was selected to select the specific proteins of *L. santarosai* serovar Alice and they were downloaded into Excel to generate results.

#### Percentage of global protein identity between the Colombian strain and a representative species of pathogenic, intermediate and saprophytic subgroups of the genus Leptospira

Percent protein sequence identity between *Leptospira* species was carried out using the RAST server. Subsequently, the presence of orthologous proteins was manually verified in the remaining species.

#### Identification and classification of the different types of CRISPR-CAS systems in L. santarosai serovar Alice

#### Reference genomes

The genomic sequences of the Colombian species of *Leptospira* from the NCBI database were used. And they were downloaded in Fasta and Genbank formats.

#### Description of Cas systems

To identify the CRISPR-Cas system and determine its class and type that were in the Colombian species, the genomes in fasta format were entered into the RAST (Rapid Annotation using Subsystem Technology) web server (22) and the CRISPRFinder software in order to identify the number of existing proteins.

#### Description of CRISPR arrays

Using the RAST web server, the Cas genes were analyzed (22), and the CRISPRDETECT server (23) was also used to finally define these arrangements.

#### Analysis of spacer sequences

The derived spacer sequences were submitted to the CRISPRTARGET database (24) to show their localization.

#### Distribution of prophage sequences and detection of phages of the Colombian strain

The genomes were downloaded in fasta format and subsequently evaluated on the PHASTER web server (25) with the aim of identifying whether genes encoding protein products corresponding to bacteriophages that are located outside the CRISPR arrays were found within the genome.

#### Biosynthesis of vitamin B12

Using the RAST program, the search was specified to a specific pattern, so the genes that had a direct synthesis function of vitamin B12 were located, as well as cofactors, transporters and coenzymes used in its synthesis. Size, function and classification into subsystems were described for each of them.

#### Data processing and analysis techniques

The analysis was done using computational biology methodologies, bioinformatics tools as described for each parameter analyzed. In addition, it was standardized with other research groups that previously used the same bioinformatic methodology even in the same bacterial genus (5), to adjust the methodology used by said groups and carried out preliminary bioinformatic analyzes with strains previously sequenced and evaluated to verify the concordance of the results.

This guaranteed optimal analysis of the genome of the strain of interest. The genome sequences of the other leptospire species that were in the databases were exhaustively analyzed to analyze their quality and those that had low quality or errors were omitted from the analyses. For the statistical analysis of the data, a descriptive analysis was used through absolute and relative frequencies.

## Results

### Molecular characterization of the Colombian strain L. santarosai serovar Alice

After total genome sequencing, it was deposited in DDBJ/EMBL/GenBank with the accession number AKWS00000000.2. Figure 1 presents the comparative phylogenetic analysis of the reference gene for classification into bacterial species with the 16s ribosomal gene and where all the *Leptospira* genomes reported until January 2020 were used. The Colombian strain was verified as *Leptospira* santarosai and a branch support value of 99% was obtained. Furthermore, it is evident in the figure that the ribosomal gene sequences of the 67 *Leptospira* species were effectively discriminated.

To delimit the phylogenetic analysis, the primers described by Merien et al. were used. 2005 (26). This region contains about 70% of the gene’s total polymorphisms. From the result of the phylogenetic analysis, the correct separation of the genus *Leptospira* with the genetically closest genus (Leptonema) was observed, and the adequate separation of the different subgroups or main clades was achieved and the ability to separate the different species with branch support was also evidenced. between 5 and 99% (Fig. 1).

It should be noted that it is not possible to completely differentiate some species from others, which states that the 16s ribosomal gene is not robust enough to individually separate the 67 species of *Leptospira*, given that the highly conserved sequence of this ribosomal gene reduces its discriminatory power and makes it difficult to identify some closely related species such as *L. meyeri and L. yanagawae or L. saintgironsiae and L. neocaledonica* among others.(2,8,17,18), although, it does manage to correctly differentiate the Colombian species *L. santarosai* serovar Alice from all other species (Fig. 1). These results are consistent with those obtained in recent studies where 65 species were analyzed (5). The above highlights the importance of investigating other markers that allow differentiating species with greater discriminatory power (5).

As evidenced by the phylogenetic analysis with the 16s ribosomal gene (Fig. 1), the classification of the 67 species into two well-defined clades is confirmed. The first and main one is subdivided into two subclades where the pathogenic and intermediate species are located. This phylogenetic classification of *Leptospira* shows that species of intermediate pathogenicity share a common ancestor with pathogenic species, and although they can be pathogenic, in experimental trials they have exhibited moderate pathogenicity both in accidental hosts and in humans and some animals and this pathogenicity It depends largely on the immune status of the host (27). The second main clade includes saprophytic species, in which species isolated from the environment and that do not cause infection are found.

In general, the analysis of the genome of the group of pathogenic leptospires shows larger size, the other groups have similar sizes. Regarding the G+C percentage, those with the highest percentage are the pathogenic and intermediate ones, which are similar, followed by the saprophytic ones. Regarding the number of tRNA genes, those with the greatest number are the pathogenic and intermediate ones, which are similar, and finally the saprophytic ones.

The Colombian strain has 37 tRNAs that represent the 20 amino acids. In contrast, the fast-growing saprophytic species L. biflexa has 35 tRNA genes (28). This finding indicates that the growth rate of this genus is not limited by a low number of tRNA genes, but is related to different metabolic capacities between different species of *Leptospira* spp (28,29). Regarding the number of CDS, they all have similar numbers and in terms of the percentage of coding, those with the highest percentage are the saprophytic ones, followed by the intermediate ones and finally the pathogenic ones. Compared to the percentage of pseudogenes, the group that presents the highest percentage is the pathogenic ones, followed by the saprophytic and intermediate ones, which have similar percentages (Table 1). This may explain the high variability and the continuous changes they can make to adapt to different hosts.

**Table 1.**
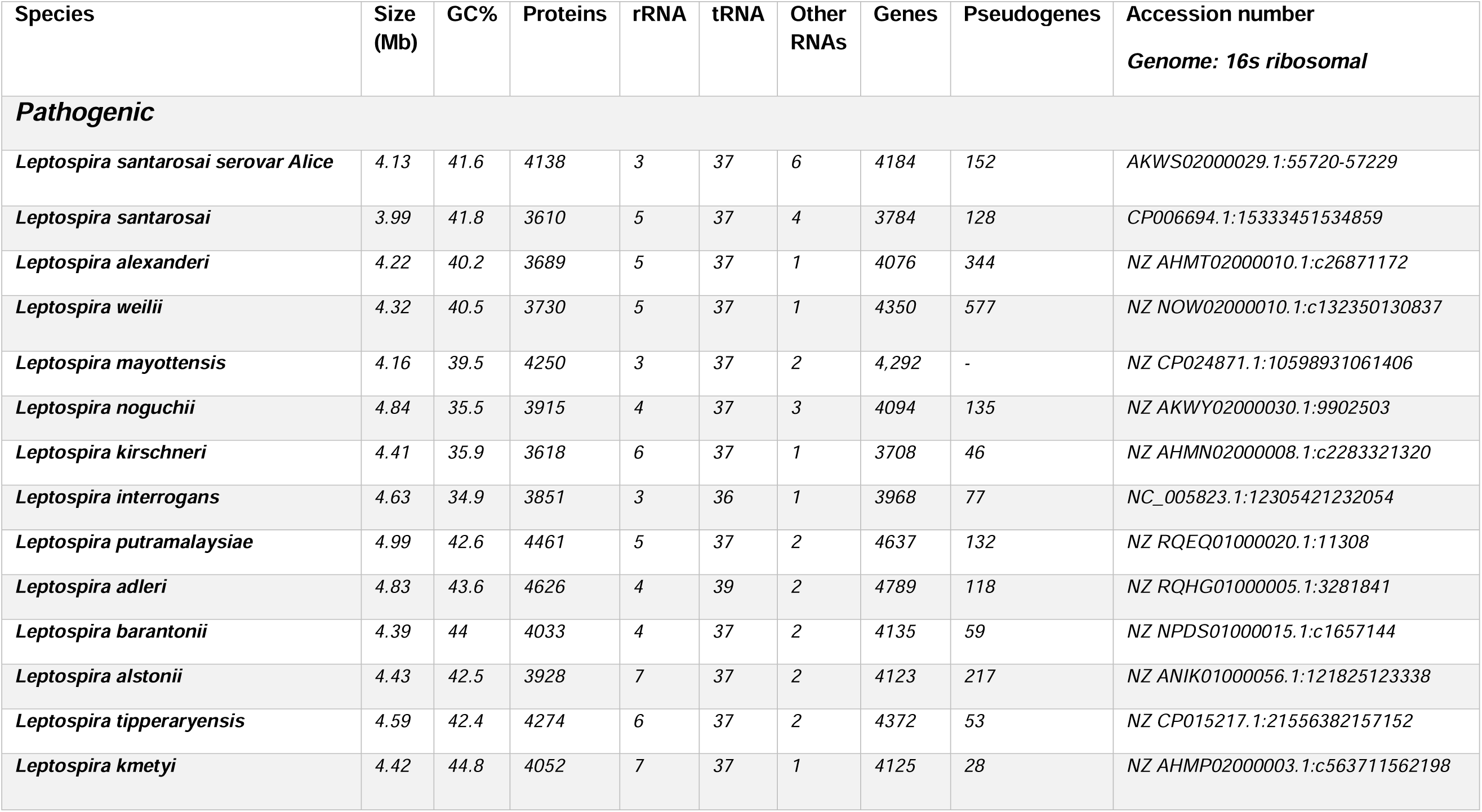
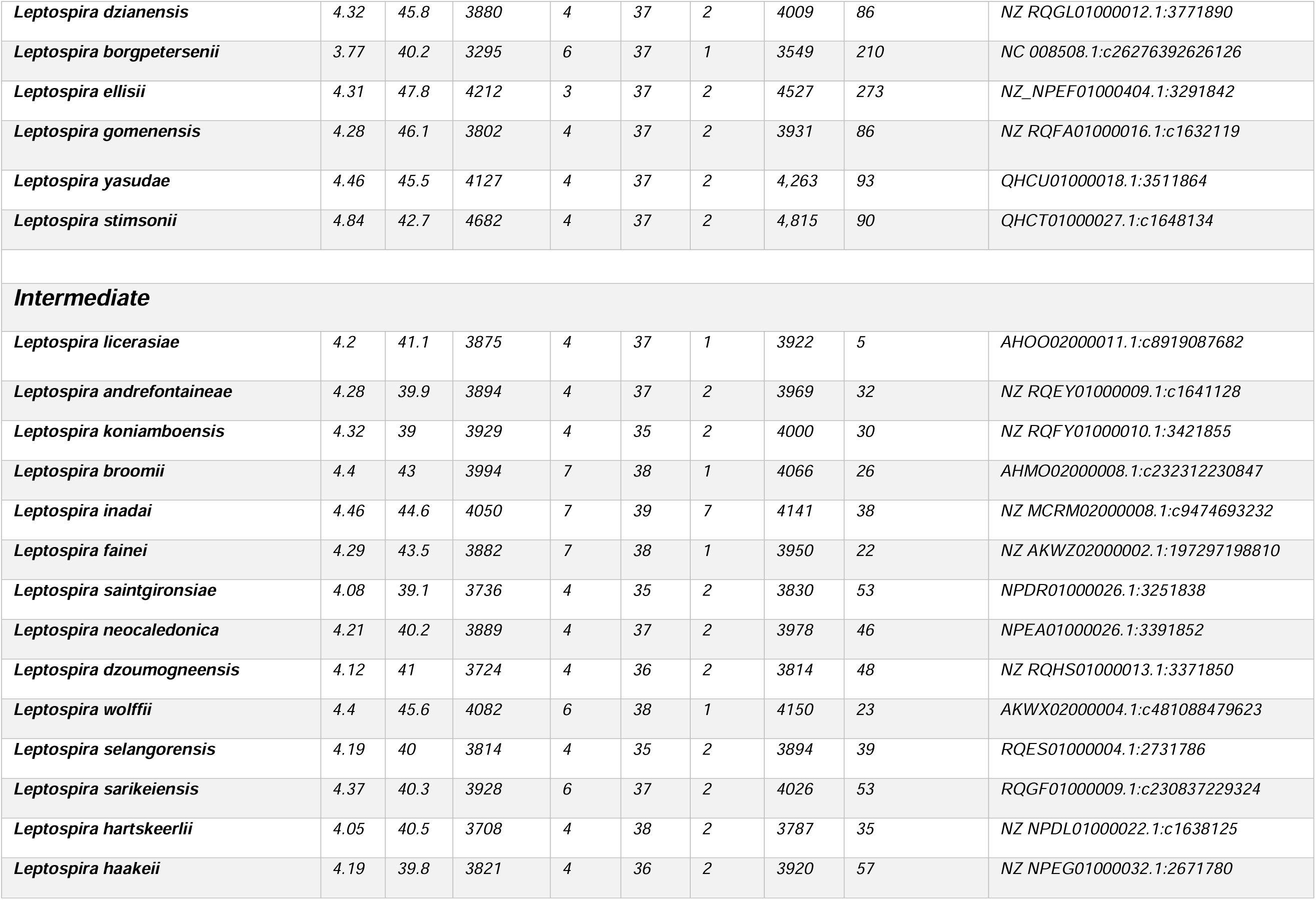
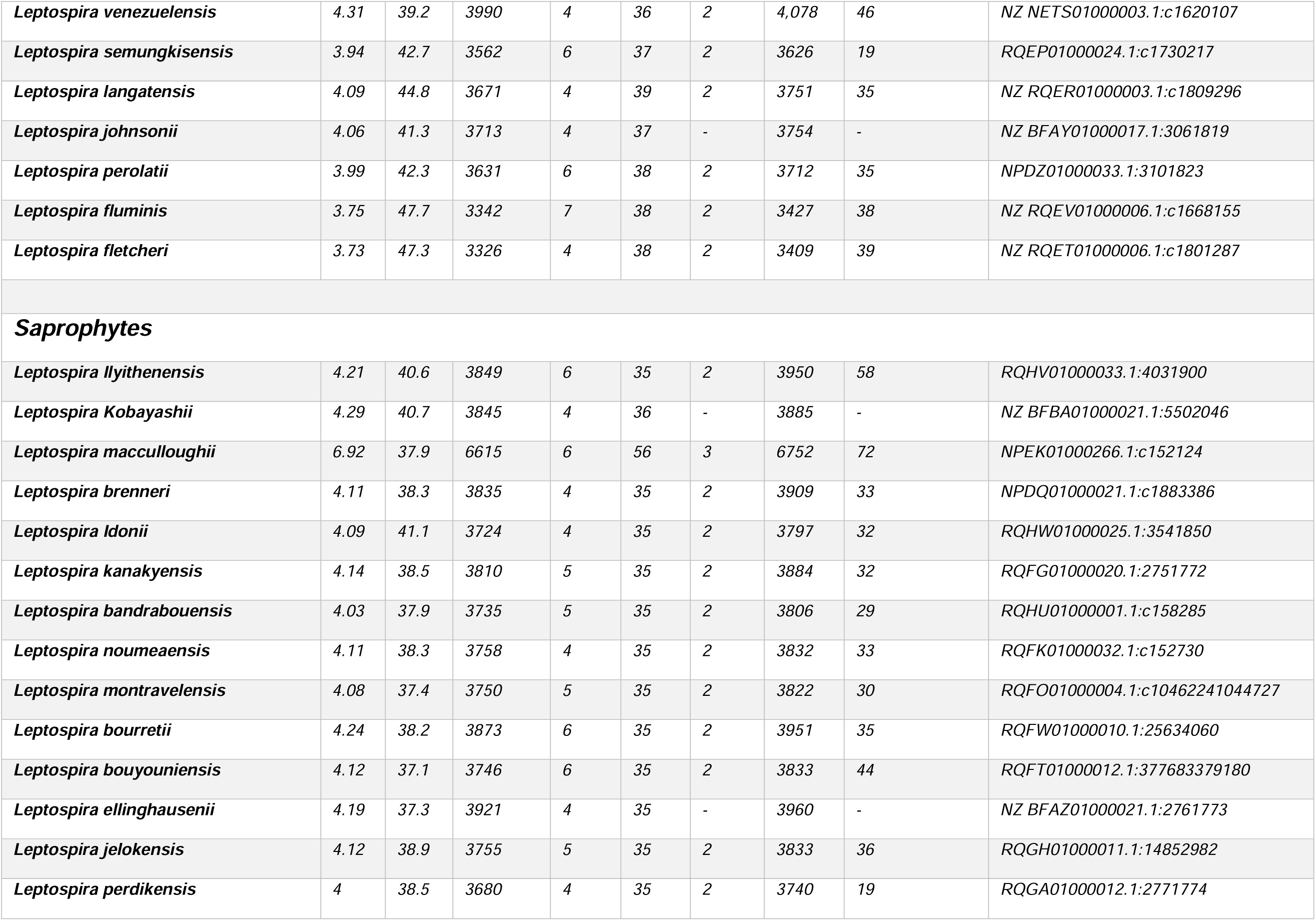
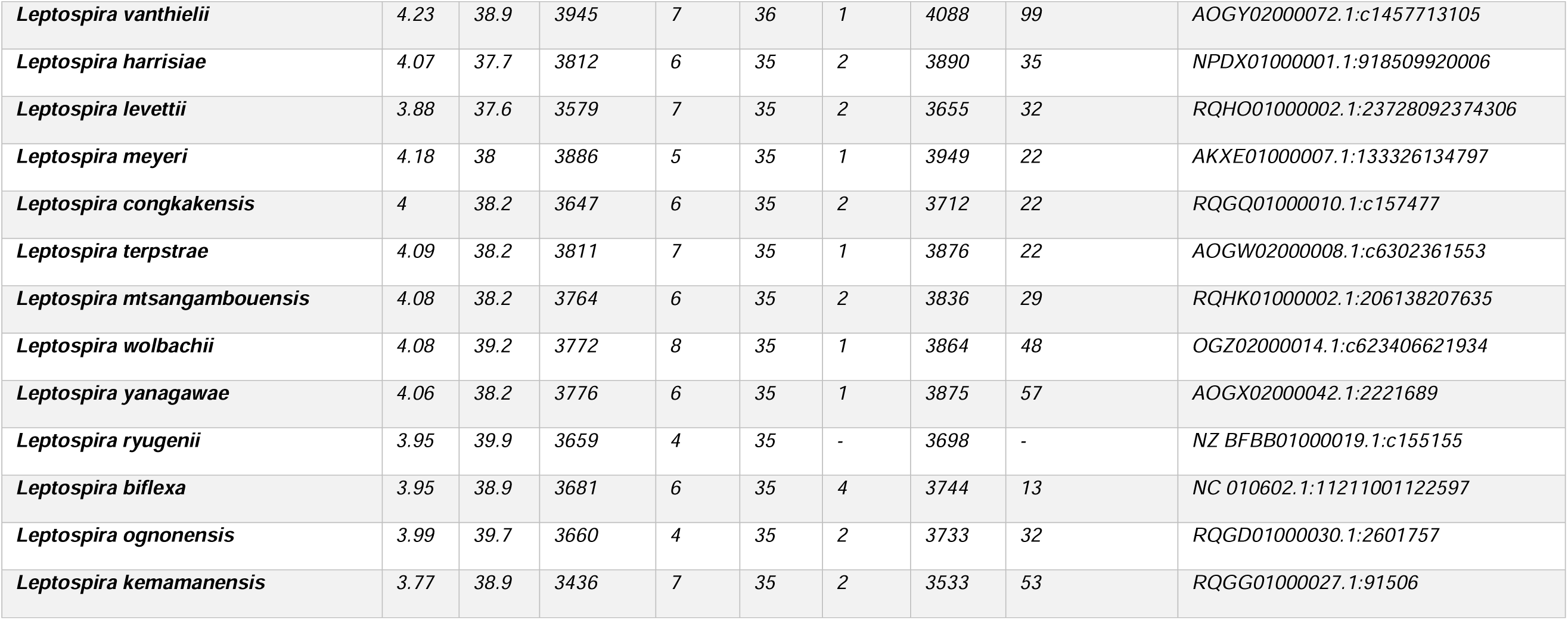
Genomic characterization of *L. santarosai* serovar Alice used in bioinformatics analyses. The genome accession number and the location of the 16s ribosomal gene within these genomes are shown, in addition to the physical characteristics of the genomes of the 67 *Leptospira* species.

| Species | Size (Mb) | GC% | Proteins | rRNA | tRNA | Other RNAs | Genes | Pseudogenes | Accession number<br>*Genome: 16s ribosomal* |
| --- | --- | --- | --- | --- | --- | --- | --- | --- | --- |
| **Pathogenic** |  |  |  |  |  |  |  |  |  |
| *Leptospira santarosai* serovar Alice | 4.13 | 41.6 | 4138 | 3 | 37 | 6 | 4184 | 152 | AKWS02000029.1:55720-57229 |
| *Leptospira santarosai* | 3.99 | 41.8 | 3610 | 5 | 37 | 4 | 3784 | 128 | CP006694.1:15333451534859 |
| *Leptospira alexanderi* | 4.22 | 40.2 | 3689 | 5 | 37 | 1 | 4076 | 344 | NZ AHMT02000010.1:c26871172 |
| *Leptospira weilii* | 4.32 | 40.5 | 3730 | 5 | 37 | 1 | 4350 | 577 | NZ NOW02000010.1:c132350130837 |
| *Leptospira mayottensis* | 4.16 | 39.5 | 4250 | 3 | 37 | 2 | 4,292 | - | NZ CP024871.1:10598931061406 |
| *Leptospira noguchii* | 4.84 | 35.5 | 3915 | 4 | 37 | 3 | 4094 | 135 | NZ AKWY02000030.1:9902503 |
| *Leptospira kirschneri* | 4.41 | 35.9 | 3618 | 6 | 37 | 1 | 3708 | 46 | NZ AHMN02000008.1:c2283321320 |
| *Leptospira interrogans* | 4.63 | 34.9 | 3851 | 3 | 36 | 1 | 3968 | 77 | NC_005823.1:12305421232054 |
| *Leptospira putramalaysiae* | 4.99 | 42.6 | 4461 | 5 | 37 | 2 | 4637 | 132 | NZ RQEQ01000020.1:11308 |
| *Leptospira adleri* | 4.83 | 43.6 | 4626 | 4 | 39 | 2 | 4789 | 118 | NZ RQHG01000005.1:3281841 |
| *Leptospira barantonii* | 4.39 | 44 | 4033 | 4 | 37 | 2 | 4135 | 59 | NZ NPDS01000015.1:c1657144 |
| *Leptospira alstonii* | 4.43 | 42.5 | 3928 | 7 | 37 | 2 | 4123 | 217 | NZ ANIK01000056.1:121825123338 |
| *Leptospira tipperaryensis* | 4.59 | 42.4 | 4274 | 6 | 37 | 2 | 4372 | 53 | NZ CP015217.1:21556382157152 |
| *Leptospira kmetyi* | 4.42 | 44.8 | 4052 | 7 | 37 | 1 | 4125 | 28 | NZ AHMP02000003.1:c563711562198 |
| *Leptospira dzianensis* | 4.32 | 45.8 | 3880 | 4 | 37 | 2 | 4009 | 86 | NZ RQGL01000012.1:3771890 |
| *Leptospira borgpetersenii* | 3.77 | 40.2 | 3295 | 6 | 37 | 1 | 3549 | 210 | NC 008508.1:c26276392626126 |
| *Leptospira ellisii* | 4.31 | 47.8 | 4212 | 3 | 37 | 2 | 4527 | 273 | NZ_NPEF01000404.1:3291842 |
| *Leptospira gomenensis* | 4.28 | 46.1 | 3802 | 4 | 37 | 2 | 3931 | 86 | NZ RQFA01000016.1:c1632119 |
| *Leptospira yasudae* | 4.46 | 45.5 | 4127 | 4 | 37 | 2 | 4,263 | 93 | QHCU01000018.1:3511864 |
| *Leptospira stimsonii* | 4.84 | 42.7 | 4682 | 4 | 37 | 2 | 4,815 | 90 | QHCT01000027.1:c1648134 |
| **Intermediate** |  |  |  |  |  |  |  |  |  |
| *Leptospira licherasiae* | 4.2 | 41.1 | 3875 | 4 | 37 | 1 | 3922 | 5 | AHOO02000011.1:c8919087682 |
| *Leptospira andrefontaineae* | 4.28 | 39.9 | 3894 | 4 | 37 | 2 | 3969 | 32 | NZ RQEY01000009.1:c1641128 |
| *Leptospira koniamboensis* | 4.32 | 39 | 3929 | 4 | 35 | 2 | 4000 | 30 | NZ RQFY01000010.1:3421855 |
| *Leptospira broomii* | 4.4 | 43 | 3994 | 7 | 38 | 1 | 4066 | 26 | AHMO02000008.1:c232312230847 |
| *Leptospira inadai* | 4.46 | 44.6 | 4050 | 7 | 39 | 7 | 4141 | 38 | NZ MCRM02000008.1:c9474693232 |
| *Leptospira fainei* | 4.29 | 43.5 | 3882 | 7 | 38 | 1 | 3950 | 22 | NZ AKWZ02000002.1:197297198810 |
| *Leptospira saintgironisae* | 4.08 | 39.1 | 3736 | 4 | 35 | 2 | 3830 | 53 | NPDR01000026.1:3251838 |
| *Leptospira neocaledonica* | 4.21 | 40.2 | 3889 | 4 | 37 | 2 | 3978 | 46 | NPEA01000026.1:3391852 |
| *Leptospira dzoumogneensis* | 4.12 | 41 | 3724 | 4 | 36 | 2 | 3814 | 48 | NZ RQHS01000013.1:3371850 |
| *Leptospira wolffii* | 4.4 | 45.6 | 4082 | 6 | 38 | 1 | 4150 | 23 | AKWX02000004.1:c481088479623 |
| *Leptospira selangorensis* | 4.19 | 40 | 3814 | 4 | 35 | 2 | 3894 | 39 | RQES01000004.1:2731786 |
| *Leptospira sarikeiensis* | 4.37 | 40.3 | 3928 | 6 | 37 | 2 | 4026 | 53 | RQGF01000009.1:c230837229324 |
| *Leptospira hartskeerlii* | 4.05 | 40.5 | 3708 | 4 | 38 | 2 | 3787 | 35 | NZ NPDL01000022.1:c1638125 |
| *Leptospira haakeii* | 4.19 | 39.8 | 3821 | 4 | 36 | 2 | 3920 | 57 | NZ NPEG01000032.1:2671780 |
| *Leptospira venezuelensis* | 4.31 | 39.2 | 3990 | 4 | 36 | 2 | 4,078 | 46 | NZ NETS01000003.1:c1620107 |
| *Leptospira semungkisensis* | 3.94 | 42.7 | 3562 | 6 | 37 | 2 | 3626 | 19 | RQEP01000024.1:c1730217 |
| *Leptospira langatensis* | 4.09 | 44.8 | 3671 | 4 | 39 | 2 | 3751 | 35 | NZ RQER01000003.1:c1809296 |
| *Leptospira johnsonii* | 4.06 | 41.3 | 3713 | 4 | 37 | - | 3754 | - | NZ BFA Y01000017.1:3061819 |
| *Leptospira perolatii* | 3.99 | 42.3 | 3631 | 6 | 38 | 2 | 3712 | 35 | NPDZ01000033.1:3101823 |
| *Leptospira fluminis* | 3.75 | 47.7 | 3342 | 7 | 38 | 2 | 3427 | 38 | NZ RQEV01000006.1:c1668155 |
| *Leptospira fletcheri* | 3.73 | 47.3 | 3326 | 4 | 38 | 2 | 3409 | 39 | NZ RQET01000006.1:c1801287 |
| **Saprophytes** |  |  |  |  |  |  |  |  |  |
| *Leptospira llyithenensis* | 4.21 | 40.6 | 3849 | 6 | 35 | 2 | 3950 | 58 | RQHV01000033.1:4031900 |
| *Leptospira Kobayashii* | 4.29 | 40.7 | 3845 | 4 | 36 | - | 3885 | - | NZ BFBA01000021.1:5502046 |
| *Leptospira macculloughii* | 6.92 | 37.9 | 6615 | 6 | 56 | 3 | 6752 | 72 | NPEK01000266.1:c152124 |
| *Leptospira brenneri* | 4.11 | 38.3 | 3835 | 4 | 35 | 2 | 3909 | 33 | NPDQ01000021.1:c1883386 |
| *Leptospira ldonii* | 4.09 | 41.1 | 3724 | 4 | 35 | 2 | 3797 | 32 | RQHW01000025.1:3541850 |
| *Leptospira kanakyensis* | 4.14 | 38.5 | 3810 | 5 | 35 | 2 | 3884 | 32 | RQFG01000020.1:2751772 |
| *Leptospira bandrabouensis* | 4.03 | 37.9 | 3735 | 5 | 35 | 2 | 3806 | 29 | RQHU01000001.1:c158285 |
| *Leptospira noumeaensis* | 4.11 | 38.3 | 3758 | 4 | 35 | 2 | 3832 | 33 | RQFK01000032.1:c152730 |
| *Leptospira montravelensis* | 4.08 | 37.4 | 3750 | 5 | 35 | 2 | 3822 | 30 | RQFO01000004.1:c10462241044727 |
| *Leptospira bourretii* | 4.24 | 38.2 | 3873 | 6 | 35 | 2 | 3951 | 35 | RQFW01000010.1:25634060 |
| *Leptospira bouyouniensis* | 4.12 | 37.1 | 3746 | 6 | 35 | 2 | 3833 | 44 | RQFT01000012.1:377683379180 |
| *Leptospira ellinghausenii* | 4.19 | 37.3 | 3921 | 4 | 35 | - | 3960 | - | NZ BFAZ01000021.1:2761773 |
| *Leptospira jelokensis* | 4.12 | 38.9 | 3755 | 5 | 35 | 2 | 3833 | 36 | RQGH01000011.1:14852982 |
| *Leptospira perdikensis* | 4 | 38.5 | 3680 | 4 | 35 | 2 | 3740 | 19 | RQGA01000012.1:2771774 |
| *Leptospira vanthielii* | 4.23 | 38.9 | 3945 | 7 | 36 | 1 | 4088 | 99 | AOGY02000072.1:c1457713105 |
| *Leptospira harrisiae* | 4.07 | 37.7 | 3812 | 6 | 35 | 2 | 3890 | 35 | NPDx01000001.1:918509920006 |
| *Leptospira levettii* | 3.88 | 37.6 | 3579 | 7 | 35 | 2 | 3655 | 32 | RQHO01000002.1:23728092374306 |
| *Leptospira meyeri* | 4.18 | 38 | 3886 | 5 | 35 | 1 | 3949 | 22 | AKXE01000007.1:133326134797 |
| *Leptospira congkakensis* | 4 | 38.2 | 3647 | 6 | 35 | 2 | 3712 | 22 | RQGQ01000010.1:c157477 |
| *Leptospira terpstrae* | 4.09 | 38.2 | 3811 | 7 | 35 | 1 | 3876 | 22 | AOGW02000008.1:c6302361553 |
| *Leptospira mtsangambouensis* | 4.08 | 38.2 | 3764 | 6 | 35 | 2 | 3836 | 29 | RQHK01000002.1:206138207635 |
| *Leptospira wolbachii* | 4.08 | 39.2 | 3772 | 8 | 35 | 1 | 3864 | 48 | OGZ02000014.1:c623406621934 |
| *Leptospira yanagawae* | 4.06 | 38.2 | 3776 | 6 | 35 | 1 | 3875 | 57 | AOGX02000042.1:2221689 |
| *Leptospira ryugenii* | 3.95 | 39.9 | 3659 | 4 | 35 | - | 3698 | - | NZ BFBB01000019.1:c155155 |
| *Leptospira biflexa* | 3.95 | 38.9 | 3681 | 6 | 35 | 4 | 3744 | 13 | NC 010602.1:11211001122597 |
| *Leptospira ognonensis* | 3.99 | 39.7 | 3660 | 4 | 35 | 2 | 3733 | 32 | RQGD01000030.1:2601757 |
| *Leptospira kemamanensis* | 3.77 | 38.9 | 3436 | 7 | 35 | 2 | 3533 | 53 | RQGG01000027.1:91506 |

#### Genome characteristics of the Colombian strain Leptospira santarosai serovar Alice

The genome of the Colombian strain *L. santarosai* consists of two circular chromosomes, the major chromosome (Chromosome I) and the minor chromosome (Chromosome II), which add up to a total of 4.13 Mb. The circular representations of both chromosomes are shown in the (Fig. 2). The two chromosomes of *L. santarosai* have a G+C percentage of 41, it has three genes for ribosomal RNA (rRNA) distributed as follows, 1 gene for the 23s subunit, 1 gene for the 16s subunit and 1 gene for subunit 5s, 6 genes of other types of RNA and in addition to 37 transfer RNA (tRNA) genes as presented in Table 1.

**Figure 2.**
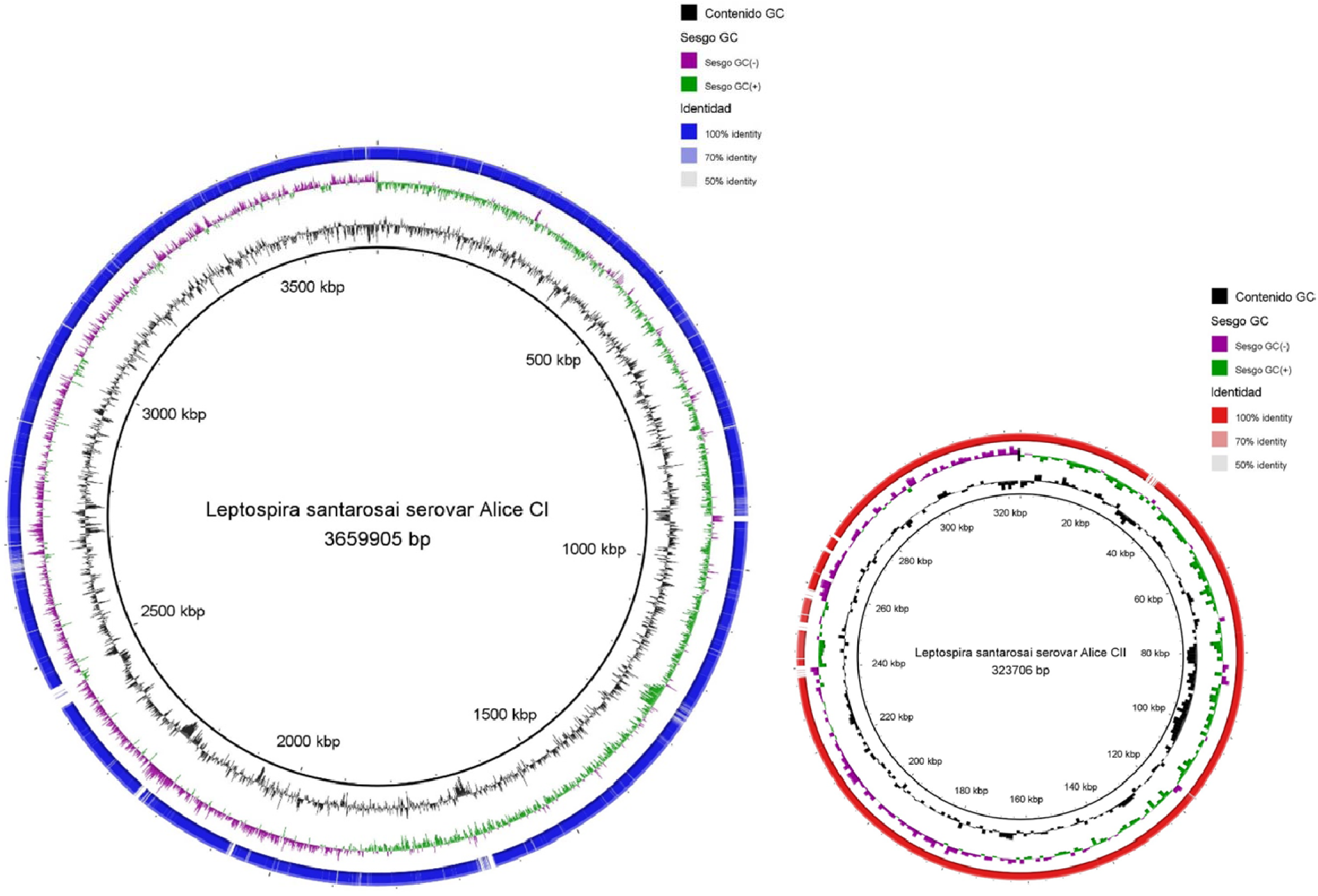
Genomic identity analysis carried out through the BRIG program between*L. santarosai serovar Shermani and the Colombian strain L. santarosai serovar Alice (thick black inner ring). The second inner ring indicates the GC content (black color) followed by the GC bias (purple and green colors). The color intensity of the outer ring blue for chromosome I and red for chromosome II represents the percent identity of nucleotides from 50 to 90.* Own elaboration by the author of this work.

#### Concatenated sequence analysis to identify serovars related to L. santarosai

22 and 23 different alleles were detected for the gyrB and secY genes, respectively. Resulting in the correct identification of 26 serovars and 3 groups with 8 indistinguishable serovars (Fig. 3).

**Figure 3.**
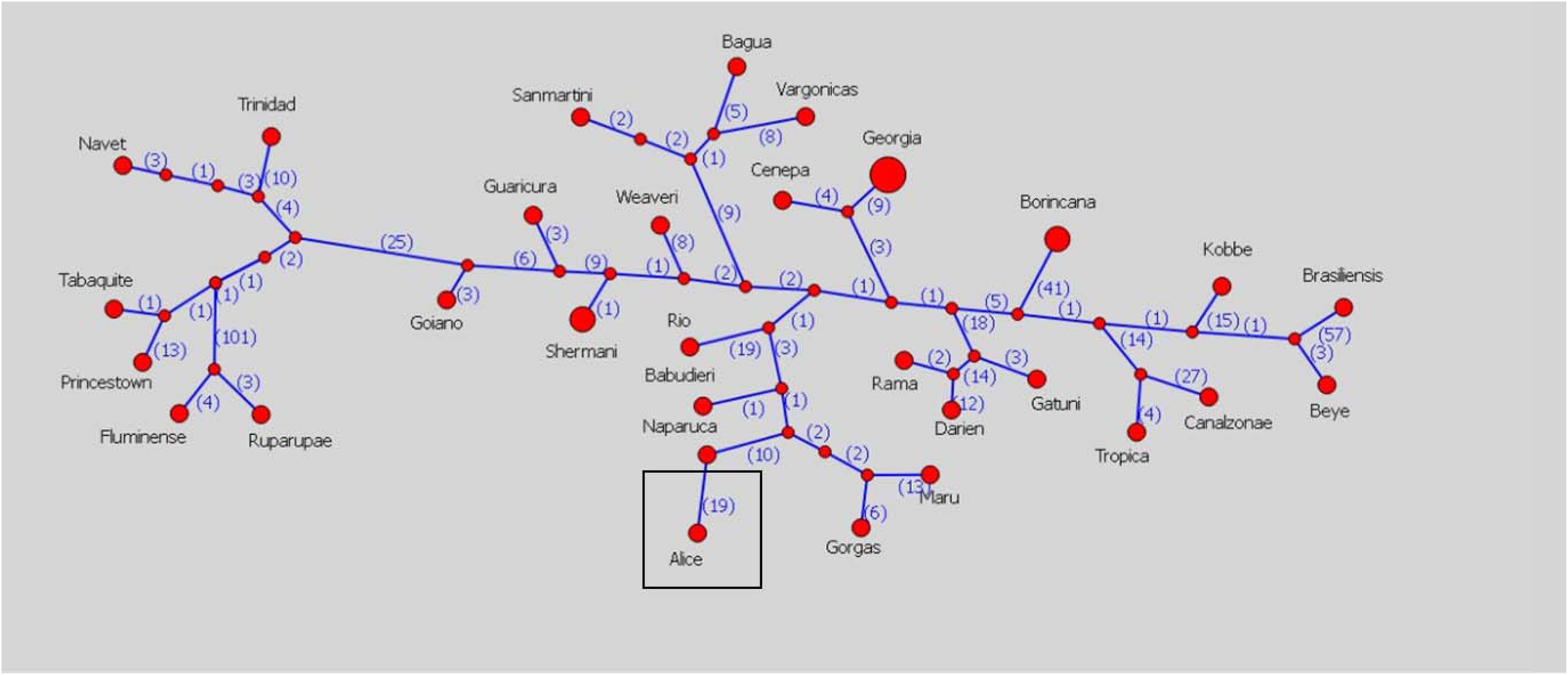
Concatenated sequence analysis for*L. santarosai*. Highly polymorphic genes are shown (*gyrB and secY) to identify serovars within the species, where serovar Alice is presented in the figure within a black square.* Own elaboration by the author of this work.

#### Analysis of genomic identity between the Colombian strain L. santarosai serovar Alice and the reference genome of L. santarosai

In the results presented in Figure 2, two chromosomal pseudomolecules were inferred for the Colombian strain, assigning proteins 3,796 and 318 to chromosomes I and II respectively. For its part, the genomic identity analysis between the Colombian strain and the reference strain resulted in a nucleotide identity of 92.95% and a high degree of synteny between the chromosomes of 95% (Fig. 4). Furthermore, 53 and 68 identical proteins were identified for serovar Shermani and the Colombian strain respectively, corresponding to 96.44% of their genome (Fig. 5). Of the 68 proteins exclusive to the Colombian strain, there are mostly uncharacterized or hypothetical proteins, then transposases, insertion elements, endonucleases, integrases, transferases and a single transmembrane protein (Table 2).

**Figure 4.**
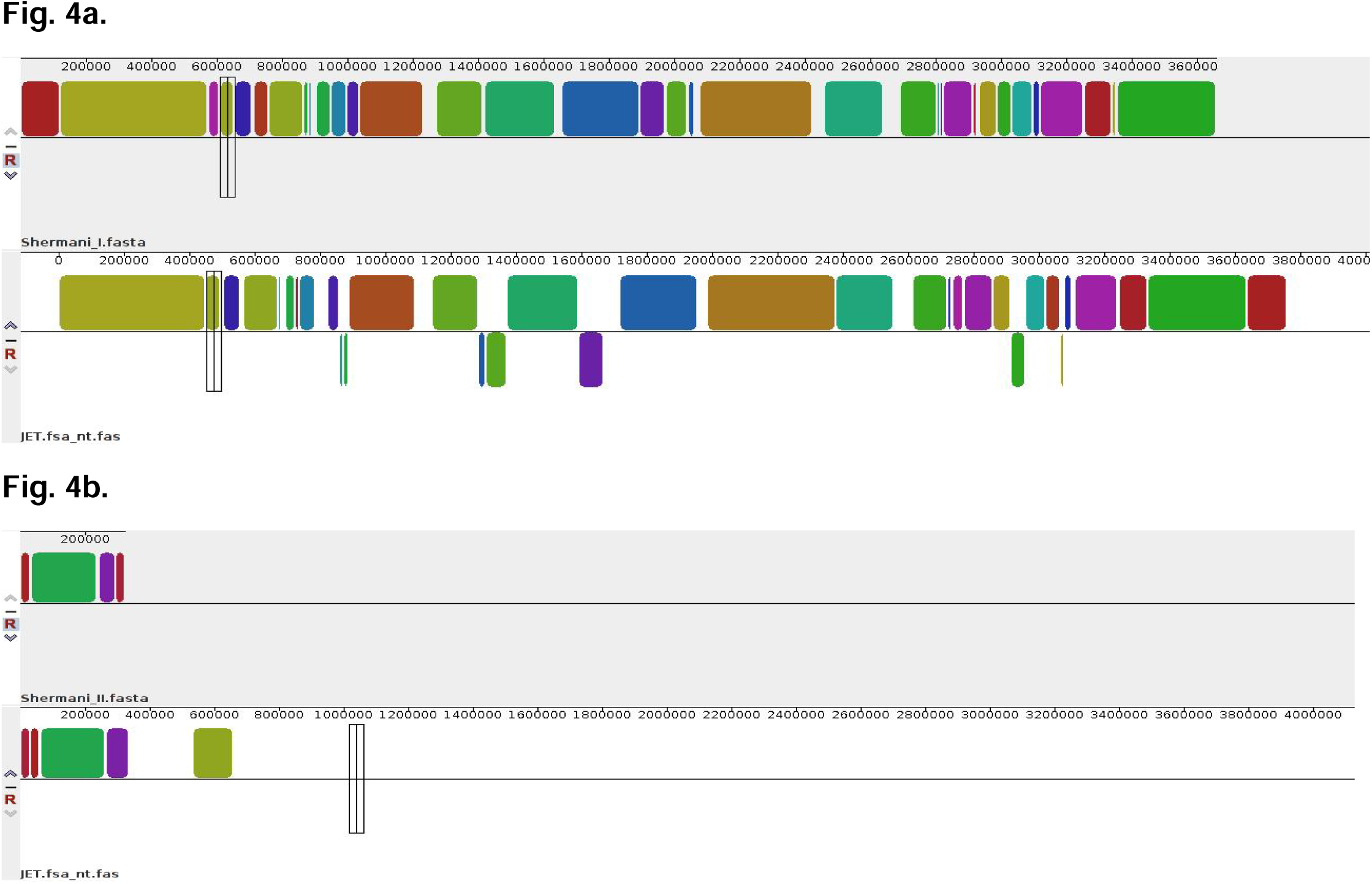
AnSynteny analysis carried out through the MAUVE program between serovars Shermani and Alice belonging to the species*L. santarosai. In part a. The analysis of chromosome I is presented and in part b. chromosome II is present. Own elaboration by the author of this work*.

**Figure 5.**
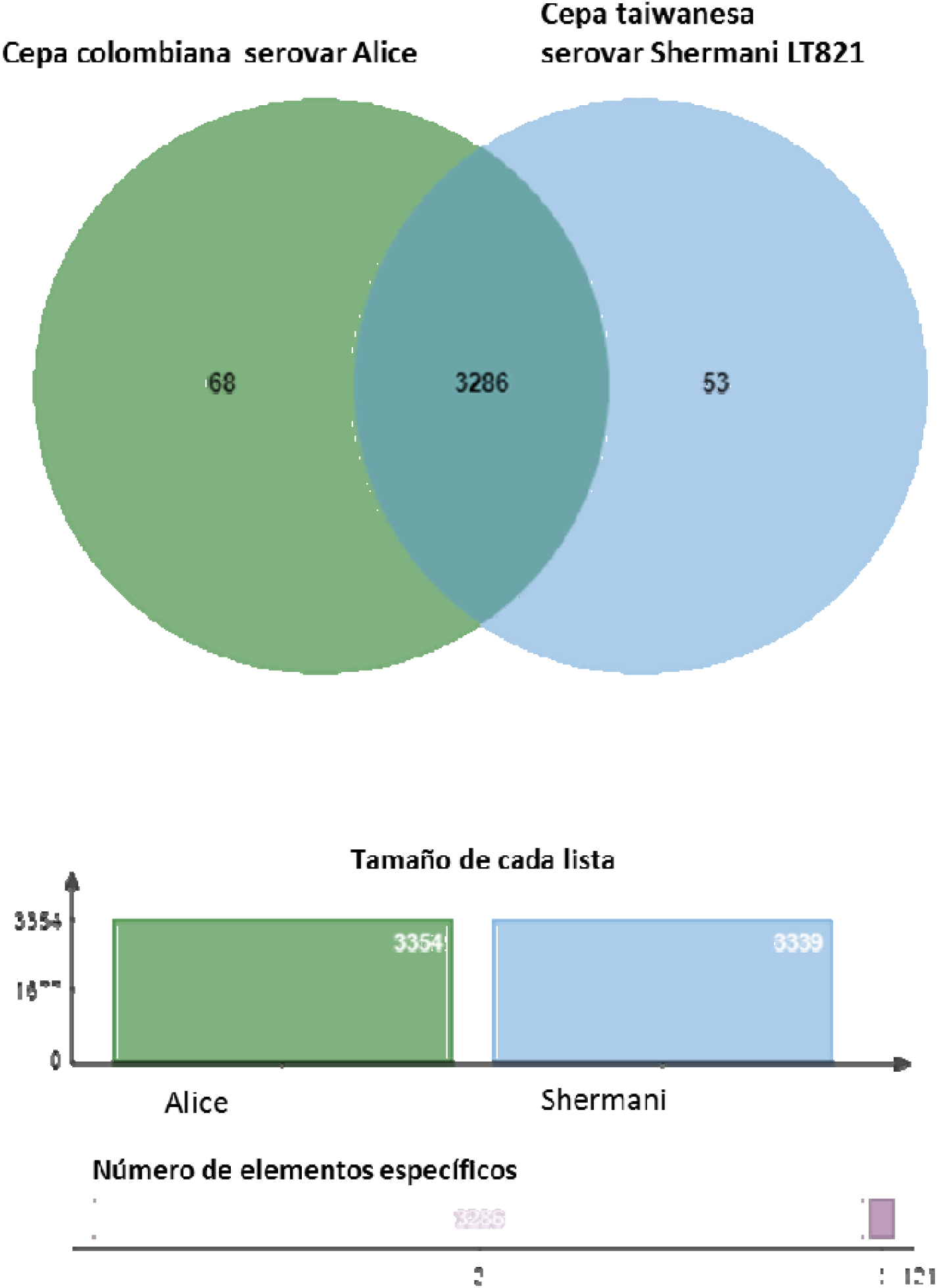
Identity analysis and detection of orthologous proteins between the species L. santarosai serovars Shermani vs Alice through the orthovenn page, for the analysis all the proteins reported for each serovar in the NCBI were downloaded in fasta format. Own elaboration by the author of this work.

**Table 2.**
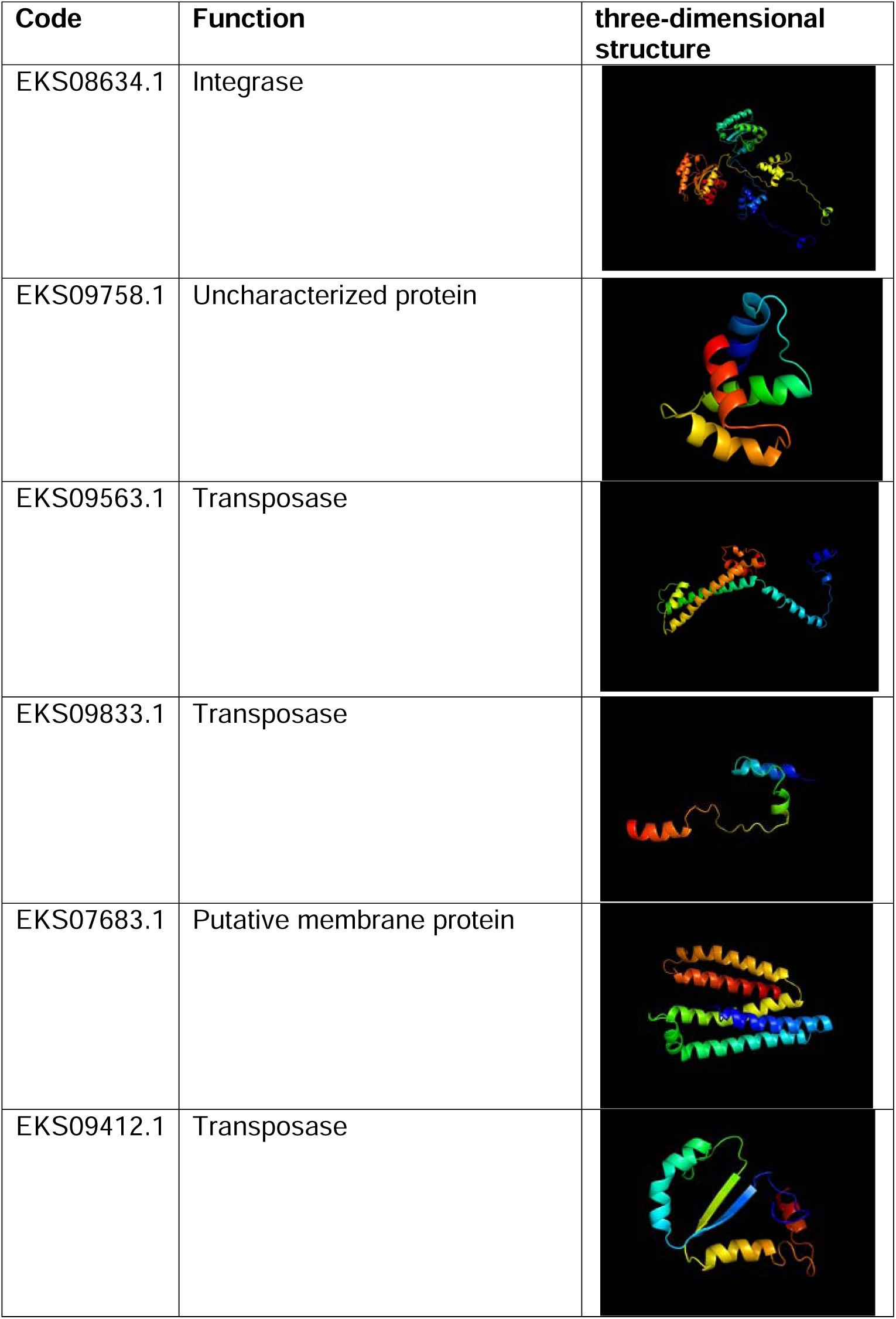
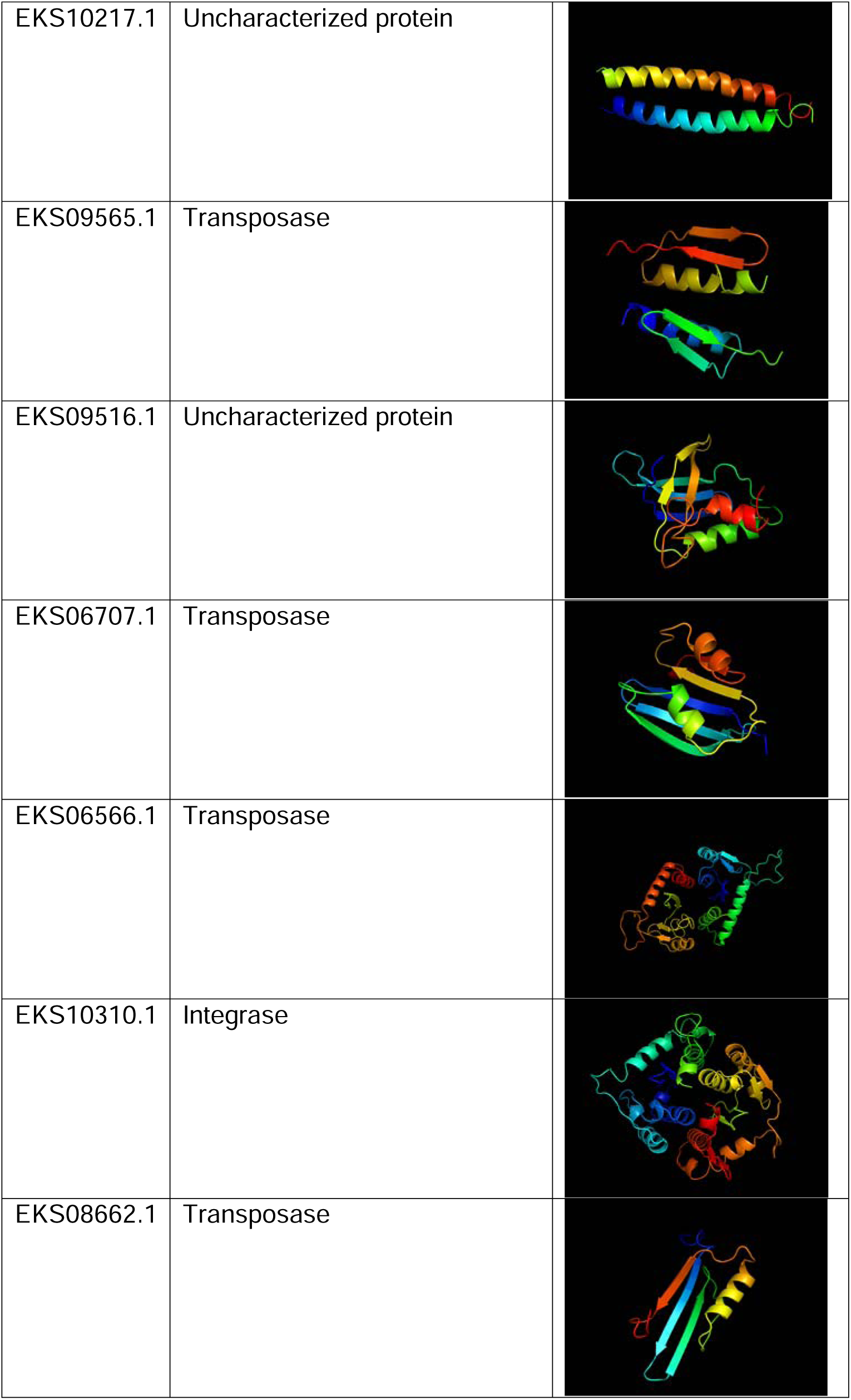
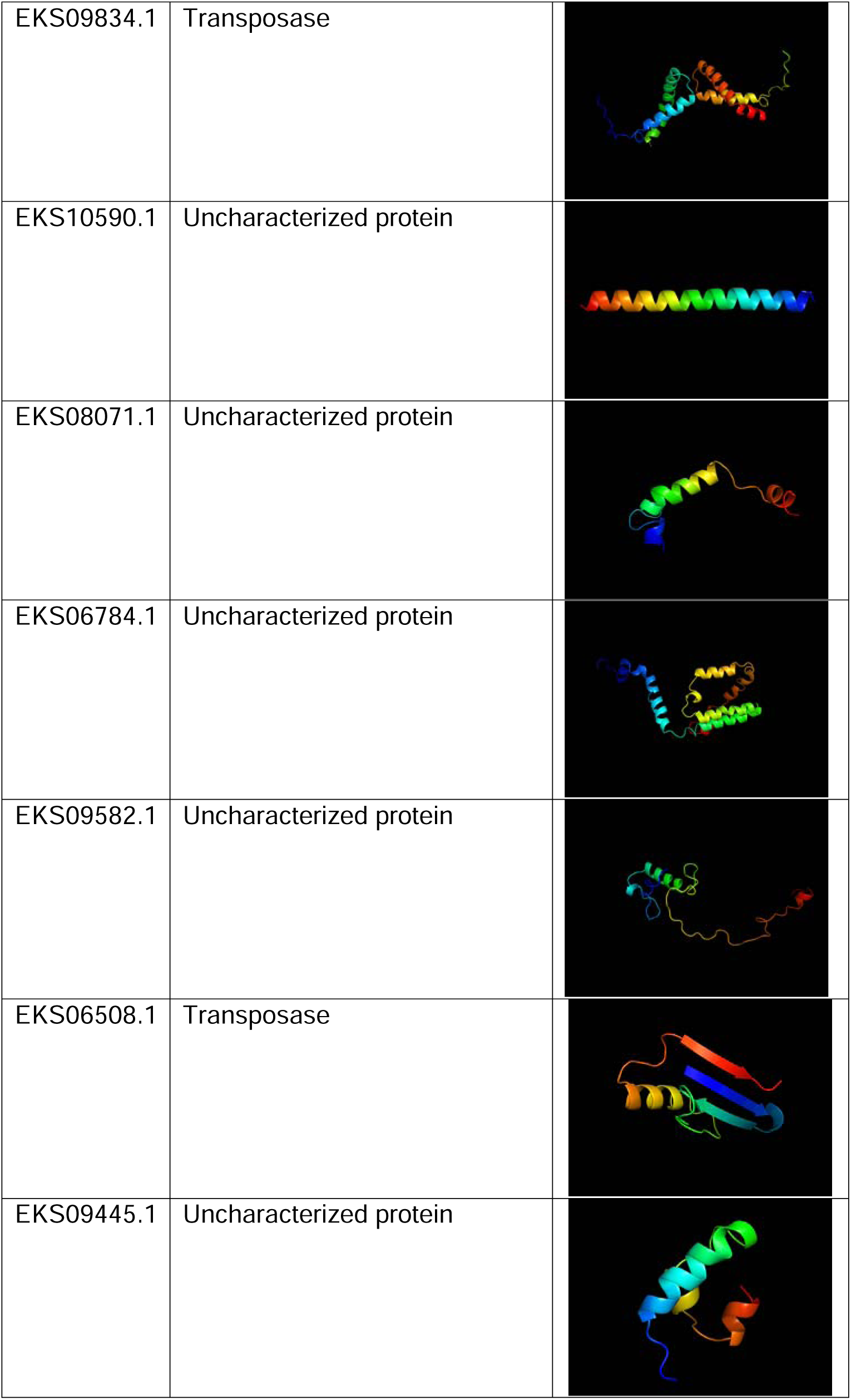
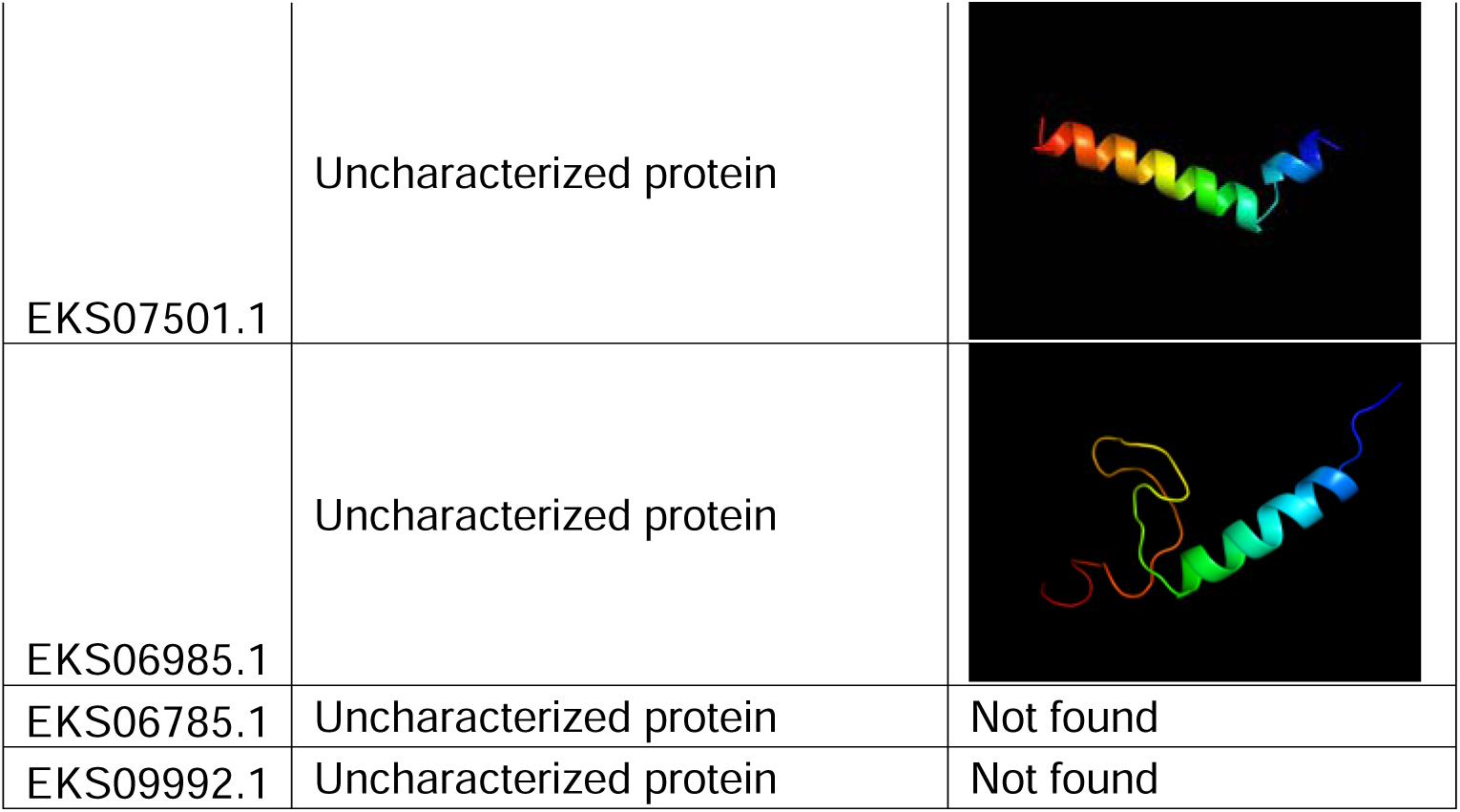
Unique proteins found in the Colombian strain *L. santarosai* serovar Alice. A total of 24 unique proteins distributed in 9 transposases, 2 integrases, 1 membrane protein and 12 proteins with no known or uncharacterized functions and their corresponding three-dimensional structures are shown.

| Code | Function | three-dimensional structure |
| --- | --- | --- |
| EKS08634.1 | Integrase |  |
| EKS09758.1 | Uncharacterized protein |  |
| EKS09563.1 | Transposase |  |
| EKS09833.1 | Transposase |  |
| EKS07683.1 | Putative membrane protein |  |
| EKS09412.1 | Transposase |  |
| EKS10217.1 | Uncharacterized protein |  |
| EKS09565.1 | Transposase |  |
| EKS09516.1 | Uncharacterized protein |  |
| EKS06707.1 | Transposase |  |
| EKS06566.1 | Transposase |  |
| EKS10310.1 | Integrase |  |
| EKS08662.1 | Transposase |  |
| EKS09834.1 | Transposase |  |
| EKS10590.1 | Uncharacterized protein |  |
| EKS08071.1 | Uncharacterized protein |  |
| EKS06784.1 | Uncharacterized protein |  |
| EKS09582.1 | Uncharacterized protein |  |
| EKS06508.1 | Transposase |  |
| EKS09445.1 | Uncharacterized protein |  |
| EKS07501.1 | Uncharacterized protein |  |
| EKS06985.1 | Uncharacterized protein |  |
| EKS06785.1 | Uncharacterized protein | Not found |
| EKS09992.1 | Uncharacterized protein | Not found |

#### Comparative genomic analysis between the Colombian strain L. santarosai serovar Alice and the genus Leptospira

The genome of the Colombian strain has approximately 4,138 protein-coding genes. According to the results obtained by the detection of orthologous proteins, a global analysis of these orthologous proteins from the 67 species of the *Leptospira* genus produced the following results, for conserved proteins: 1,650 specific to the *Leptospira* genus (Fig. 6), which which constitutes its genomic core or central genome, 409 specific to the pathogenic group, 680 specific to the intermediate group and 144 specific to the saprophytic group. Within the subgroup of pathogenic species, the Colombian strain shares the highest number of orthologous proteins with *L. santarosai* serovar Shermani 3,286 and the lowest number of orthologous proteins with L. kemamanensis 1,864. Furthermore, the Colombian strain shares orthologous proteins with members of the different subgroups of species that range between 3,286-2,600 with the pathogenic species, 2,416-2,296 with the intermediate species and 1,972-1,864 with the saprophytic species (Fig. 7). It is evident that the Colombian strain presents a greater number of its orthologous proteins with the pathogenic species (Fig. 8), which demonstrates a strong genetic relationship between the Colombian strain and this subgroup.

**Figure 6.**
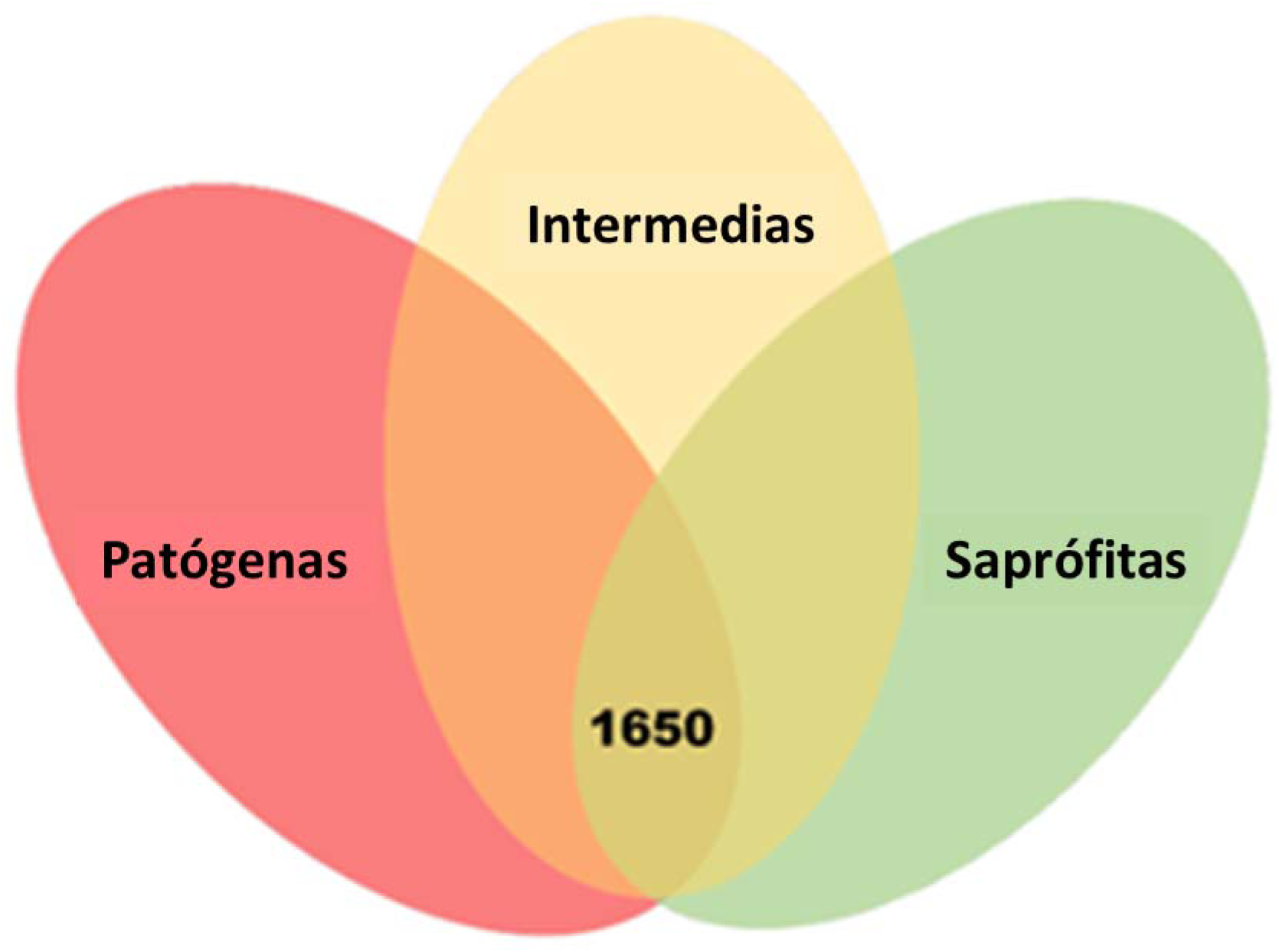
Detection of genomic core through the distribution of orthologous proteins among 67 species of the genus *Leptospira*. The red, yellow and green circles represent the pathogenic, intermediate and saprophytic subgroups. The number of orthologous proteins that are conserved in the *Leptospira* genus is highlighted. Detection of orthologous proteins was performed using a combination of OrthoVenn, OrthoMCL, and InParanoid bioinformatics tools. Own elaboration by the author of this work.

**Figure 7.**
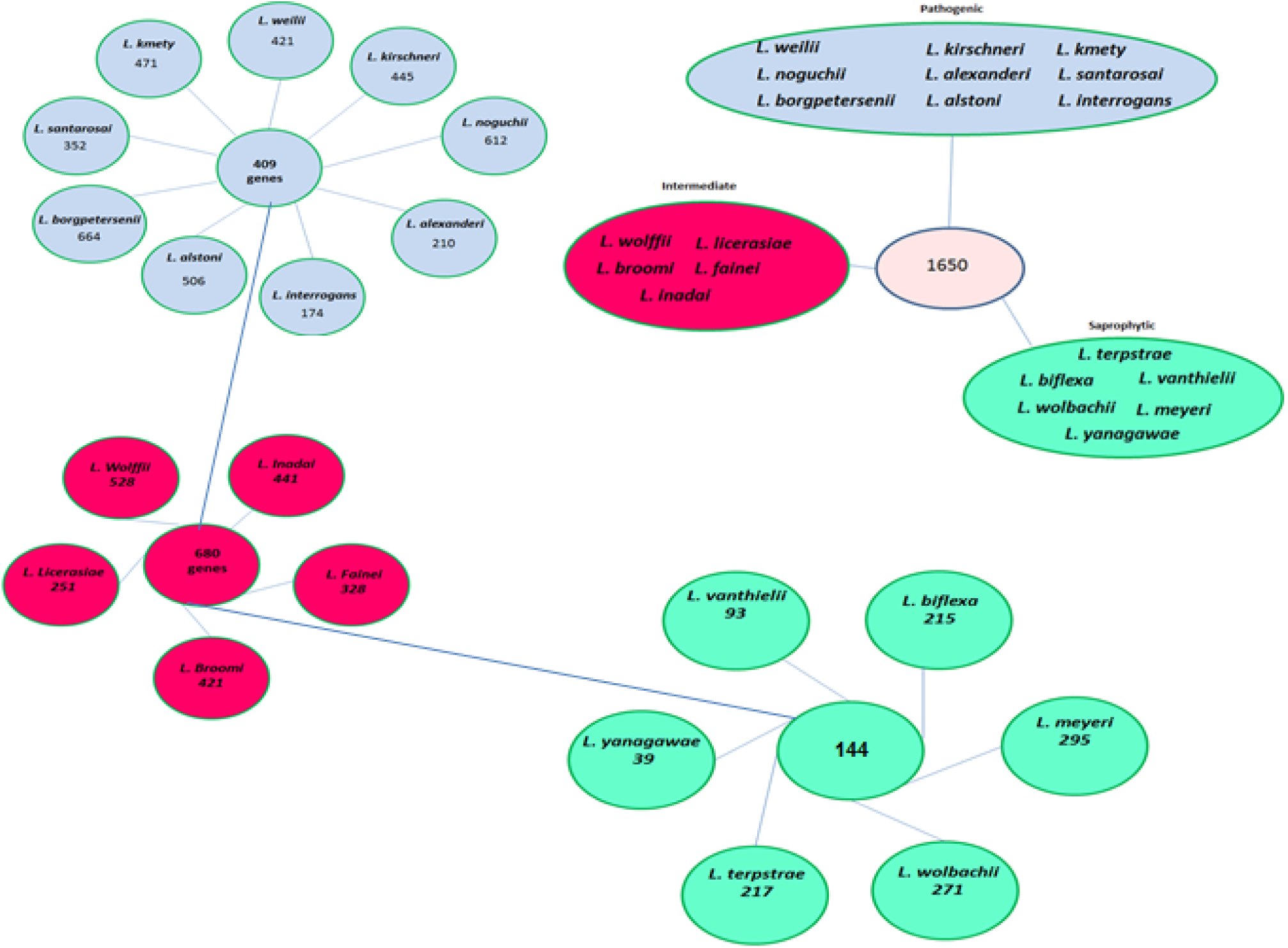
Identification of orthologous proteins of the genus*Leptospira spp. Orthologous proteins among 20 species of the genus Leptospira are represented through a combination of the OrthoMCL and InParanoid algorithms. Own elaboration by the author of this work*.

**Figure 8.**
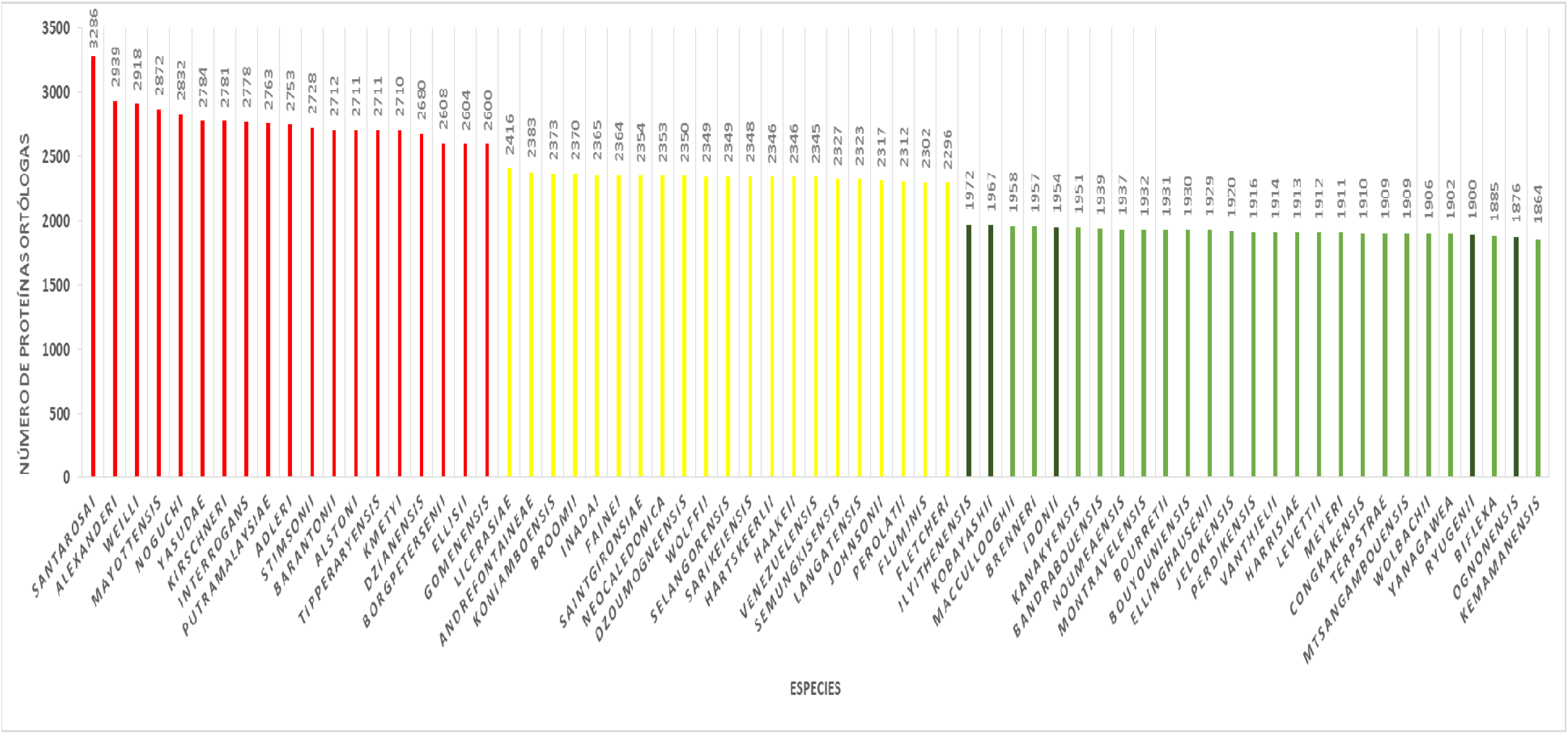
Presentation of the set of orthologous proteins detected between*L. santarosai*serovar Alice and the 67 species of the genus*Leptospira individually. The species reported until December 2020 were used, pathogenic species are shown in red, intermediate species are shown in yellow, and saprophytic species are shown in green. The number of orthologous proteins from zero to 3,500 is shown on the Y axis and all Leptospira species are presented on the X axis.*Each analysis was performed separately on the OrthoVenn server and then pooled. Own elaboration by the author of this work.

In addition, 24 unique proteins were identified (Table 2) from the Colombian species and when compared with the species representing the three groups of the bacterial genus, nine of which are transposases that participate in the transposition of insertion sequences, two integrases, one protein membrane and 12 hypothetical proteins with no known function (Fig. 10,13). Through the detection of orthologous proteins between serovars and isolates related to L. santarosai, it was possible to demonstrate great genetic diversity even among isolates that belong to the same species, it was evident that the greatest similarity in terms of orthologous proteins in their species was with the AIM strain with 3,491 orthologous proteins (Fig. 9).

**Figure 9.**
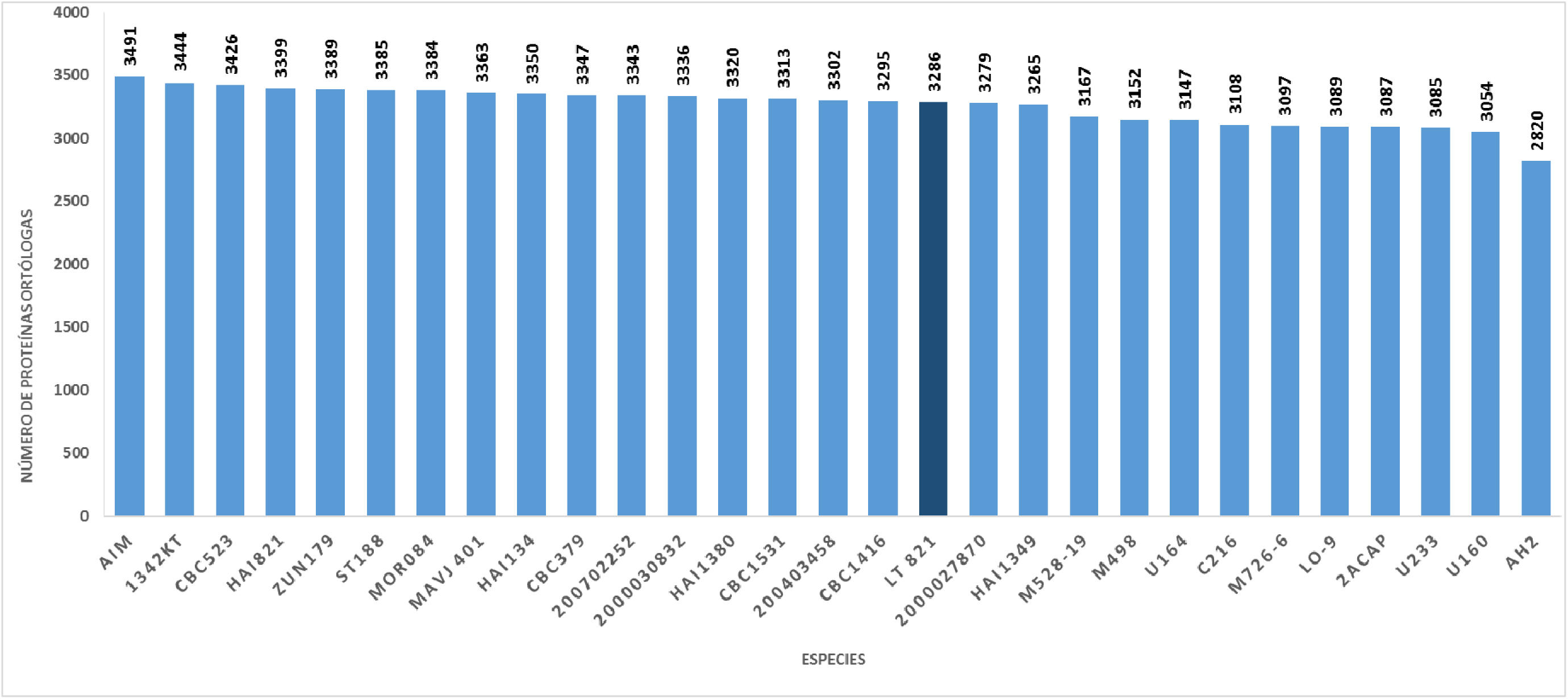
Set of orthologous proteins detected between*L. santarosai*serovar Alice and all 29 strains of the species*L. santarosai described worldwide; The genome of L. santarosai serovar Shermani strain LT 821 is highlighted in dark blue, of which its genome is complete and it is the world reference strain in this species.*Each analysis was performed separately on the OrthoVenn server and then pooled. Own elaboration by the author of this work.

**Figure 10.**
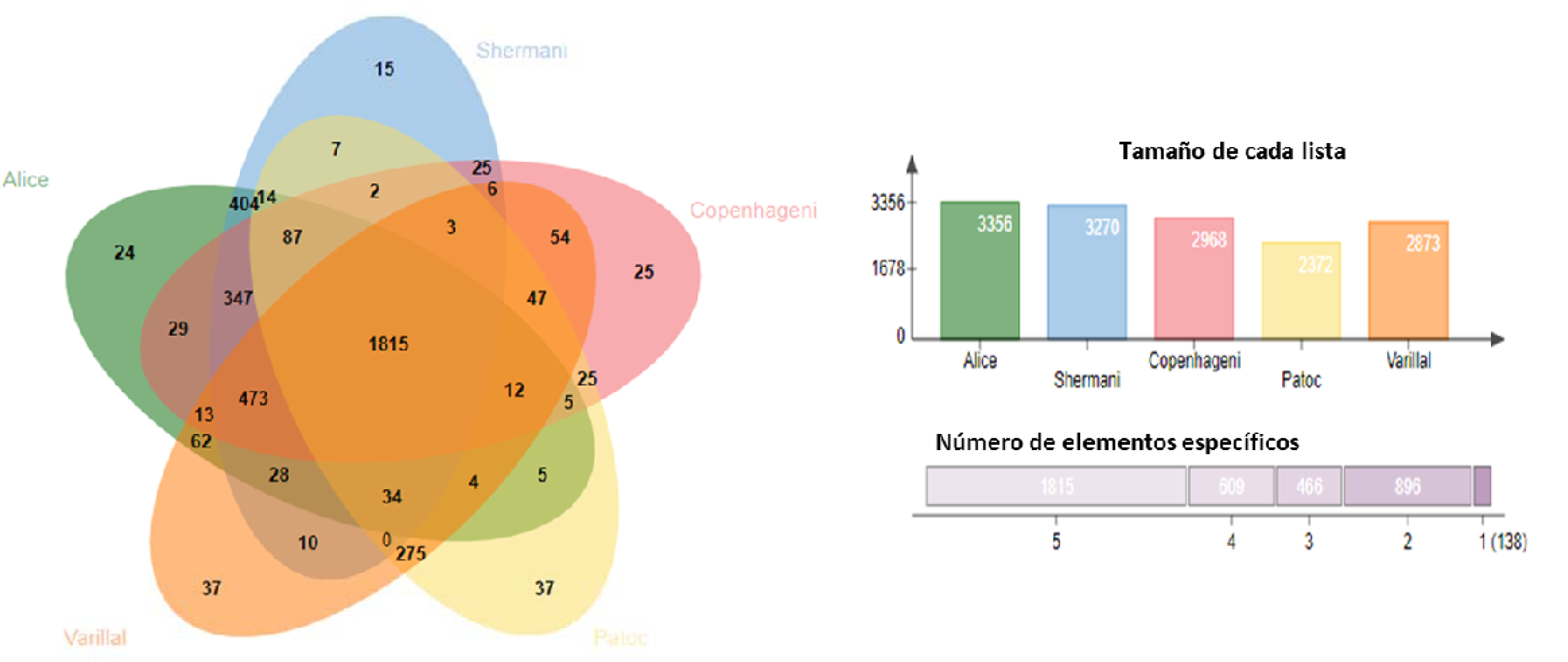
Detection of the pangenome with the PanOCT and Orthovenn software showing the common orthologous proteins of its core genome and non-common proteins of its variable genome and, for the analysis, the species were used*L. interrogans serovar Copenhageni (pathogenic), L. licerasiae serovar Varillal (Intermedia), L. biflexa serovar Patoc (Saprophytic), L. santarosai*serovar Shermani (pathogenic)*and L. santarosai serovar Alice (pathogenic). All proteins reported for each species and serovar in the NCBI database were downloaded in fasta format.* Own elaboration by the author of this work.

When analyzing the pangenome, with the PanOCT tool and representing it with Orthovenn software, it is evident that the *Leptospira* genus has an open pangenome (Fig. 10) and increases with the inclusion of new sequenced strains. It should be noted that pathogenic species contain the most open pangenome and to date have the largest number of unique genes among the different subgroups and this number is increasing given that in pathogenic species alone, the pangenome is close to 6,252 genes.

The results obtained with the GGsearch bioinformatics tool indicated that 40.98% of the proteins present in the genome of the Colombian strain have a high percentage of identity with the proteins of *L. interrogans* (80-100% similarity), while that 59.02% of the proteins are more similar to intermediate and saprophytic species with a percentage of identity ranging from 1 to 80% (Table 3).

**Table 3.**
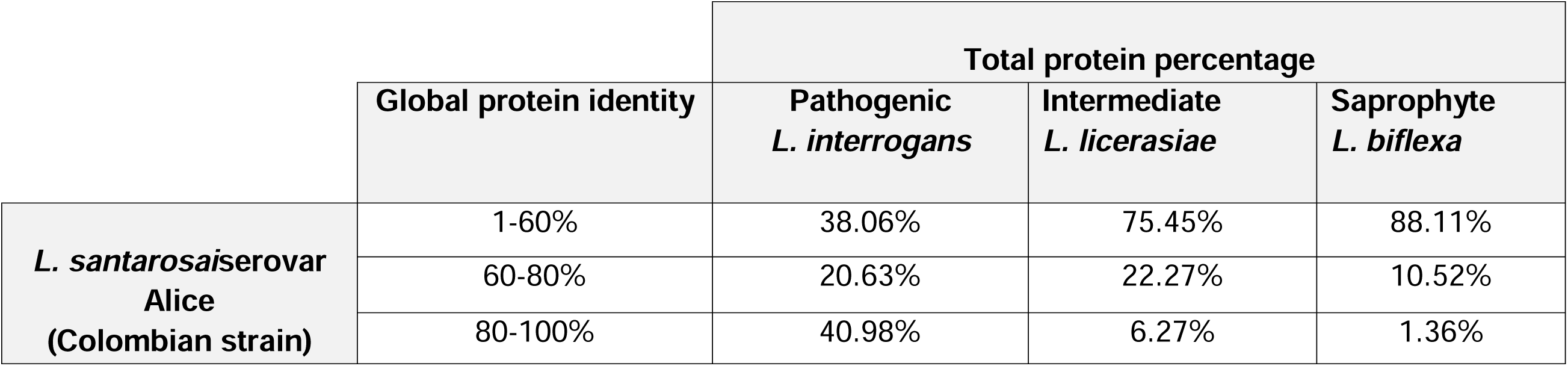
Global protein similarity or identity between the Colombian strain*L. santarosai*serovar Aliceand a representative species of the genus*Leptospira among pathogenic (L. interrogans), intermediate (L. licerasiae) and saprophytic (L. biflexa) subgroups*.

|  | Global protein identity | Total protein percentage |  |  |
| --- | --- | --- | --- | --- |
|  |  | Pathogenic<br>*L. interrogans* | Intermediate<br>*L. licerasiae* | Saprophyte<br>*L. biflexa* |
| ***L. santarosaisero* var Alice**<br>**(Colombian strain)** | 1-60% | 38.06% | 75.45% | 88.11% |
|  | 60-80% | 20.63% | 22.27% | 10.52% |
|  | 80-100% | 40.98% | 6.27% | 1.36% |

Similarly, when searching for the percentage of identity (Fig. 11) using the Seed-viewer program, similar results were found, where it is evident that the Colombian species has a percentage of identity greater than 95% with the pathogenic strain L. interrogans, while the other species representing the intermediate and saprophytic subgroups reach percentages of 50 and 10% respectively. This reaffirms the global genomic similarity of the Colombian strain with the pathogenic strains.

**Figure 11.**
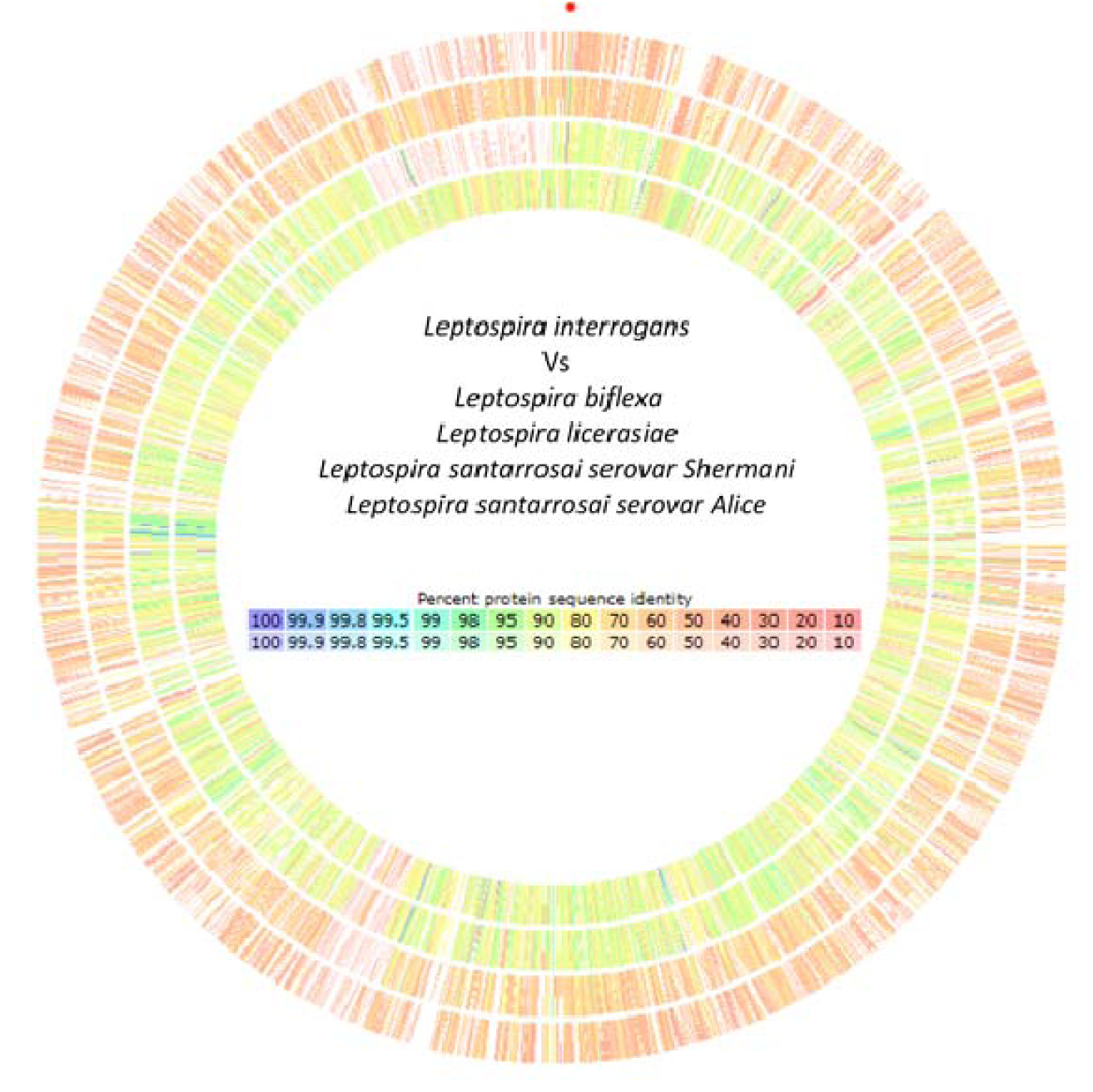
Percent identity of protein sequences between different groups of Leptospira species. As a reference genome. The species Leptospira interrogans (pathogenic) was taken. From the outside to the inside the order corresponds to Leptospira biflexa (saprophytic), Leptospira licerasiae (intermediate), Leptospira santarosai (pathogenic) and Leptospira santarosai serovar Alice (Colombian pathogenic species). Genomes stored in NCBI were used as input files. The Seed-viewer program was used for the analyses. Own elaboration by the author of this work.

#### CRISPR/Cas systems in the Colombian species

In this study, the classification criteria of the CRISPR-Cas system were followed, which divide them into two varieties (Class 1 and Class 2). The CRISPR-Cas class 1 system (CRISPR1) is made up of three types (I, III and IV), while the class 2 system (CRISPR2) comprises two types (II and V), with subdivision into 16 different subtypes according to the different Cas protein combinations (23).

When analyzing the genome of the Colombian strain of *Leptospira* santarosai, it was found that they contained CRISPR/Cas repeats and 9 families belonging to class 1, type I and subtype IA and IE were located where each of these subtypes contained 8 cas genes (Table 4) and (Fig. 12).

**Figure 12.**
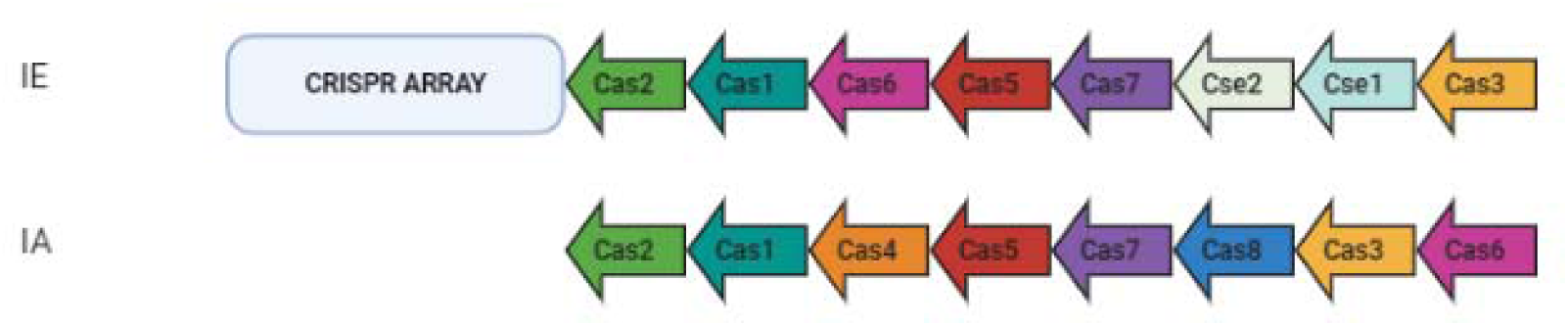
Representation of the CAS loci of the CRISPR-CAS system of the Colombian strain *L. santarosai* serovar Alice. Each of the Cas proteins is symbolized by an arrow of different colors that indicates the direction of transcription and the different CRISPR arrays. The two subtypes found, IE and IA, are represented.Own elaboration by the author of this work.

**Figure 13.**
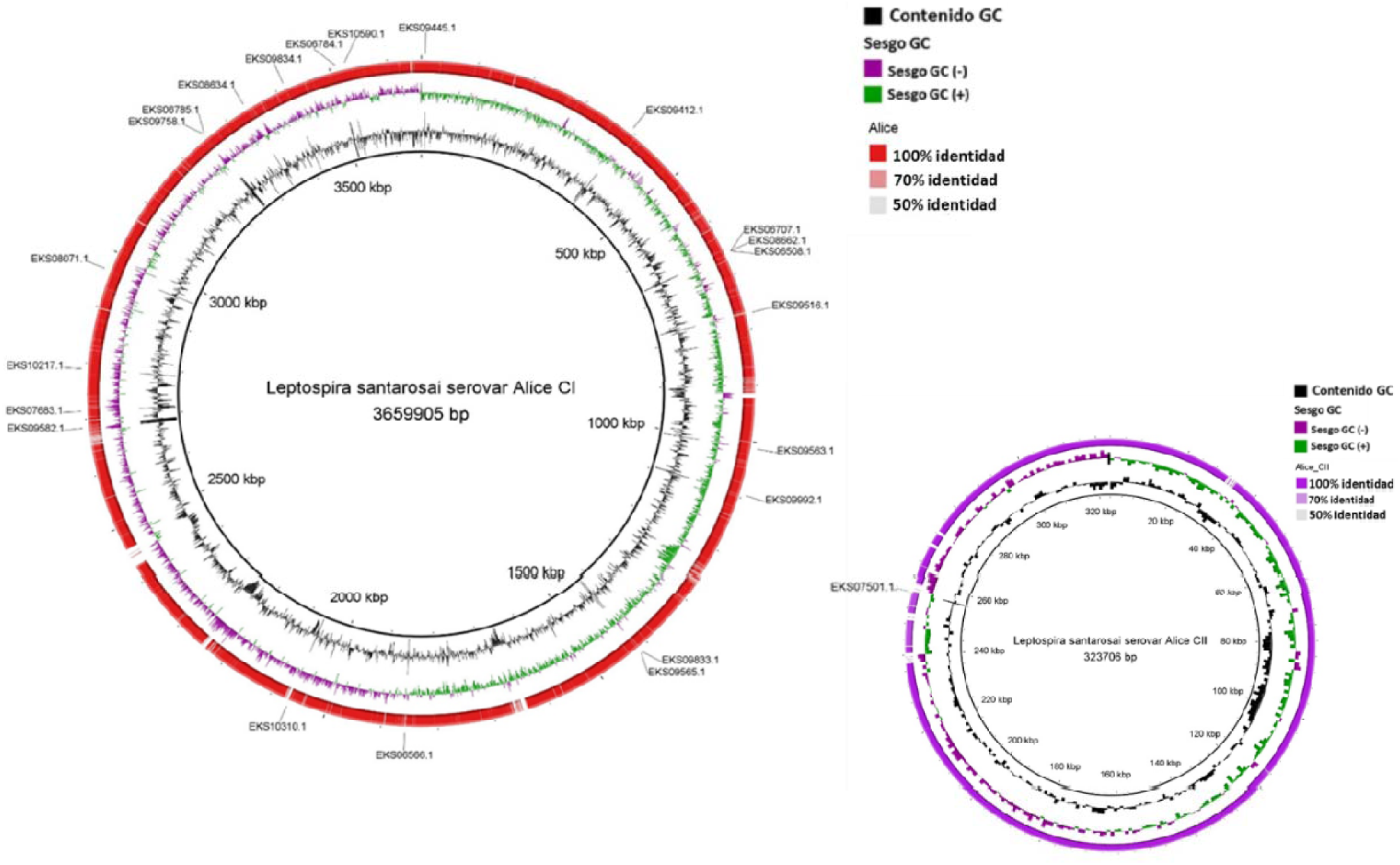
Circular map of the location of 24 unique proteins of the Colombian species. Once these proteins were detected by the PanOCT software, their search and subsequent location within the chromosome was carried out. The comparative sequence analysis was carried out with BRIG using as a reference. This result was visualized at the chromosome level through the BRIG program, using the species *Leptospira* santarosai serovar Shermani strain LT 821 as a reference chromosome and comparing it with the genome. from *Leptospira* santarosai serovar Alice (thick black inner ring). The second inner ring indicates the GC content (black color) followed by the GC bias (purple and green colors). The outermost labels show the location of the genes encoding proteins unique to the Colombian species. The intensity of the color of the red ring for chromosome I and purple for chromosome II represents the percentage of nucleotide identity from 50 to 90. Own elaboration by the author of this work.

**Table 4.**
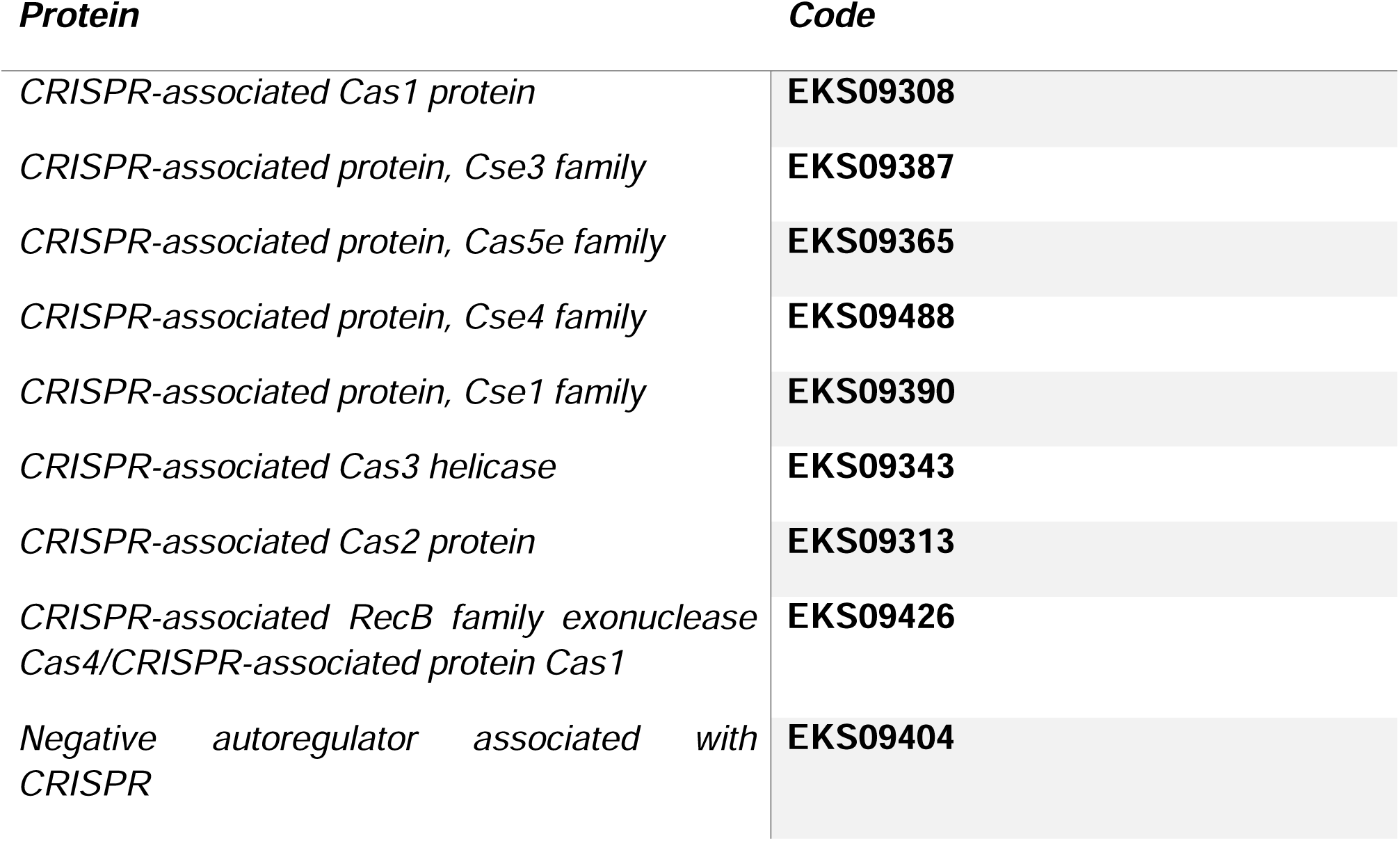
CRISPR-associated proteins in the Colombian species *L. santarosai* serovar Alice. The proteins of interest detected using the RAST server are presented. The protein and the NCBI access code are named for each family.

#### Biosynthesis of vitamin B12

In the Colombian strain, a total of 25 proteins were found generally involved in processes related to vitamin B12 (Table 5). These proteins can be divided into subsystems such as coenzymes necessary for the biosynthesis of the vitamin, cofactor transporters, acquisition of the vitamin from the external environment through specific receptors, synthesis and excretion of cobalamin. Of the total of 25 proteins found, it was observed that 15 were directly involved in the de novo biosynthesis of vitamin B12, verified through their function, demonstrating cobalamin (B12) autotrophy as a bacterial virulence factor.

**Table 5.** Proteins involved in the biosynthesis of vitamin B12 in the Colombian strain L. santarosai.

| **Protein** | **Code** |
| --- | --- |
| *Sirohydrochlorin cobaltochelatase CbiX (long) / Putative 2Fe-2S ferredoxin CbiW involved in B12 biosynthesis* | NZ_AKWS02000039 |
| *FIG056361: hypothetical protein involved in the biosynthesis of coenzyme B12* | NZ_AKWS02000039 |
| *ABC transporter, substrate binding protein (group 8, B12/iron complex)* | NZ_AKWS02000045 |
| *B12 binding domain/kinase domain/methylmalonyl-CoA mutase* | NZ_AKWS02000050 |
| *BtuB outer membrane vitamin B12 receptor* | NZ_AKWS02000077 |
| *Class II ribonucleotide reductase (coenzyme B12 dependent)* | NZ_AKWS02000079 |
| *Cobalt-zinc-cadmium resistance protein CzcD* | NZ_AKWS02000001 |
| *Magnesium-cobalt transport protein CorA* | NZ_AKWS02000021 |
| *Cobalt-zinc-cadmium resistance protein CzcA; CusA cationic efflux system protein* | NZ_AKWS02000035 |
| *Sirohydrochlorin cobaltochelatase CbiX (long) / Putative 2Fe-2S ferredoxin CbiW involved in B12 biosynthesis* | NZ_AKWS02000039 |
| *Cobalt-precorrin-5B(C1)-methyltransferase* | NZ_AKWS02000039 |
| *Cobalt-precorrin-8 methylmutase* | NZ_AKWS02000039 |
| *Cobalt-precorrin-7 (C5)-methyltransferase / Cobalt-precorrin-6B C15-methyltransferase [decarboxylating] (EC 2.1.1.196)* | NZ_AKWS02000039 |
| *Cobalt-precorrin-2 C(20)-methyltransferase* | NZ_AKWS02000039 |
| *Cobalt-precorrin 5A hydrolase* | NZ_AKWS02000039 |
| *Cobalt-precorrin-3 C(17)-methyltransferase/Cobalt-factor III reductase* | NZ_AKWS02000039 |
| *Cobalt-precorrin-4 C(11)-methyltransferase (EC 2.1.1.271)* | NZ_AKWS02000039 |
| *Cob(I)alamine adenosyltransferase Cob(I)alamine adenosyltransferase, grouped with cobalamin synthesis* | NZ_AKWS02000039 |
| *Hypothetical CobX protein related to cobalamin biosynthesis* | NZ_AKWS02000039 |
| *Cobalt-zinc-cadmium resistance protein CzcA; CusA cationic efflux system protein* | NZ_AKWS02000058 |
| *Cobalt-zinc-cadmium resistance protein CzcA; CusA cationic efflux system protein* | NZ_AKWS02000058 |
| *Cobalamin synthase* | NZ_AKWS02000060 |
| *Cobalt/zinc/cadmium efflux transporter RND, membrane fusion protein, CzcB family* | NZ_AKWS02000062 |
| *Cobalt-zinc-cadmium resistance protein CzcD* | NZ_AKWS02000063 |
| *Cobalt-zinc-cadmium resistance protein CzcA; CusA cationic efflux system protein* | NZ_AKWS02000077 |

#### Phage system in the Colombian species

In the Colombian strain, three species of different phage structures were found that could facilitate infection and subsequent colonization in the host. They are the basal plate of the phage (access code EKS06898) that serves as a binding support for the proteins of the phage tail fibers (access code EKS08862). These are extensions that are essential in the absorption stage, where It serves as the initial attachment of the phage to the host cells in order to subsequently introduce its genetic material and TMP-Phage tail measuring protein (access code EKS10262) which regulates the length of the phage tail and is located in the internal tail canal (Table 6).

**Table 6.** Detection of phages of the Colombian strain L. santarosai. The proteins that form phage structures and the access codes are presented.

| *Protein* | *Code* |
| --- | --- |
| phage basal plate | EKS06898 |
| Phage tail fiber proteins | EKS08862 |
| TMP-Phage tail measuring protein | EKS10262 |

## Discussion

Colombia is considered an endemic country for leptospirosis, with an annual increase in the number of leptospirosis cases reported since 2007 (31). Currently, *L. santarosai* is recognized as an important causative agent of human and animal leptospirosis in Colombia, causing clinical manifestations ranging from mild forms to severe complications (12). Similar results have been found on the island of Taiwan, where the main causative agent of leptospirosis is *L. santarosai* serovar Shermani. This serovar has been associated with different clinical forms of the disease, ranging from mild forms to severe clinical complications such as Weil’s disease and severe pulmonary hemorrhage syndrome without the presence of jaundice (32,33), reflecting a behavioral similar of *L. santarosai* in different geographical regions. This species has been associated with at least 56 different serovars (34); These serovars have been reported to cause infection in humans and animals (12). Furthermore, the detection of some serovars has been reported in environmental water sources (35), reflecting the ubiquity of this species. But despite this *L. santarosai* belongs to the least studied species of pathogenic leptospires.

Due to the presence in Colombia of *L. santarosai* and some serovars related to this species, a complete comparative genomic analysis is required, in addition to the implementation of molecular tools for adequate identification. *L. santarosai* has a reference genome and 29 different sequenced isolates, two of them belonging to Colombian strains (12). Therefore, given the epidemiological importance of this species in Colombia, a comparative genomic analysis contributes to discovering new molecular targets to identify species of the genus *Leptospira* and serovars related to L. santarosai. In the current research, we present a comparative genomic analysis between a Colombian strain and the entire *Leptospira* genus.

Leptospirosis is known to cause a potentially fatal pathology in humans, but the processes involved in the response to pathogenic species of this genus are not fully understood. Therefore, the genomic analyzes shown in this study allow a better understanding of the evolution of species, especially the Colombian species.

The Colombian strain is one of two Colombian *Leptospira* isolates that have been sequenced at the genomic level (32). Therefore, this genome was chosen to perform a comparative genomic analysis, because it is a strain native to Colombia that had not been reported circulating in Colombia and because the preliminary results obtained in a recent investigation showed that its use as a biological model was important., which helped clarify hypotheses from previous results where its virulence condition was widely demonstrated (3).

The Colombian strain was sequenced and characterized by phylogenetic classification using the 16s ribosomal gene, which is a global reference target for taxonomic analysis. This is the first phylogenetic classification carried out with the 67 species using the 16s ribosomal gene, the previous work carried out by Vincent AT et al. 2019, where the identification of the Colombian species previously carried out (5) is confirmed, managed to make a classification with the species that were submitted in the NCBI databases until that moment. This classification resulted in the Colombian species belonging to the pathogenic species L. santarosai.

By analyzing the classification using the complete phylogenetic tree of the 67 species of *Leptospira*, it is suggested that all species evolved from the same common ancestor, which subsequently separated into two main well-defined clades, a pathogenic clade and another saprophytic one, where the first main clade contains the pathogenic and intermediate species and the second main clade contains the saprophytic species. From the first main clade it can be suggested that all pathogenic species could have evolved from the same environmental ancestor through the acquisition of new functions through genetic transfers associated with adaptation to new hosts.

Something interesting that was validated in previous studies and that is related in this work at least at a phylogenetic level with the 16s ribosomal gene, is that within the pathogenic clade, several subgroups have been defined based on their virulence (result in patients and / or virulence in the animal model) and phylogenomic analysis (5). Thus, the subgroup that contains the Colombian species *L. santarosai* serovar Alice, L. santarosai, L. interrogans, L. kirschneri, L. noguchii, L. borgpetersenii, L. mayottensis, L. alexanderi and L. weilii are related to most frequently with serious and fatal infections in humans. These species diverged from the pathogenic clade at a specific node of evolution and may have evident closeness at the species level or at least contain common virulence factors that produce fatal outcomes with some regularity (5).

Classification based on Bayesian analysis of ribosomal 16s gene sequences has been generally accepted worldwide for *Leptospira*, especially for the classification of pathogens from non-pathogens (potential contaminants). However, the 16s ribosomal gene is highly conserved so the species in the phylogenetic tree cannot be subdivided further than shown in the figure, as they lack different characteristics within their sequences that are sufficient to make a robust phylogenetic derivation. at the species level (4.5).

For this reason, this gene does not achieve total differentiation of all the 67 species of *Leptospira*, which is why it does not allow us to distinguish some species from others, such as L. macculloughii from L. levettii. Therefore, the phylogenetic analysis of the present study allows us to represent the diversity of this bacterial genus, but it highlights the shortcomings of this gene to make a strong and exact phylogenetic inference that allows separating all species specifically.

Vincent AT et al. 2019 (5) proposes the reconstruction of the speciation process using the Supertree approach (most robust taxonomic tree – super tree), classified 65 species of *Leptospira* spp into two main groups of pathogenic and saprophytic where it locates a subclade related to saprophytic species but not It is still well discriminated and is composed of *Leptospira* idonii and four new species *L. kobayashii, L. ilyithenensis, L. ryugenii and L. ognonensis*. In this phylogenetic reorganization, pathogenic species were divided into subgroups P1 and P2 (pathogenic and intermediate respectively), in addition it is also observed that the other main clade, which is the saprophyte, is subdivided into two well-defined subclades, so a new classification was proposed, where this second main clade contains the two subgroups called S1, which includes the classic saprophytic species and a new clade S2, with the new saprophytic species (5). Therefore, it should be noted that the 16s ribosomal gene does not allow us to differentiate this new S2 clade suggested by Vincent AT et al. 2019 (5), since it was impossible to separate clades S1 from S2 proposed by this author, in fact, some species of clade S2 such as *L. ilyithenensis, L. kobayashii, L. idonii, L. ryugenii and L. ognonensis*, are found dispersed throughout the saprophytic clade without showing evolutionary closeness typical of this subgroup. The phylogenetic analyzes presented in this study allow a better understanding of the evolution of the species that form the different clades and subclades. Despite the limitations of using a single gene and this 16s gene in particular to achieve robust phylogenetic classification.

The different genomic characteristics of the pathogenic, intermediate and saprophytic subgroups generally show differences in size since the pathogenic ones have a larger size, G+C% where the intermediate ones have a larger %, number of genes where the pathogenic and intermediate ones have a greater number, transfer RNA, another type of RNA where the intermediate ones have a greater number and pseudogenes where the pathogenic ones have the greatest number, which shows the enormous variability and evolution of this genus and among the different subgroups (Table 1).

*Leptospira santarosai*It is found in several countries around the world, and contains several serovars that include several hosts in addition to different environmental sources. In the present work, once the genome of *L. santarosai* serogroup Autumnalis serovar Alice was sequenced, circular maps of the major and minor chromosomes of this bacterium were generated and comparisons were made with other genomes of pathogenic, intermediate and saprophytic species to search for orthologs. Analysis of the sequences, domains and structures thereof was used to elucidate the functions of the proteins and description of hypothetical proteins, finding notable similarities with respect to the pathogenic strains.

In the two pseudochromosomes for the Colombian strain, a high symmetric identity was found between genomes (92.95%) of serovars Shermani and the Colombian strain serovar Alice, which reflects a high degree of genetic relationship between both. Furthermore, 3,286 orthologous proteins were found when these two strains of the same species were compared.

Given the close relationship between some serovars and their hosts, the identification of serovars is epidemiologically important for the implementation of adequate prevention and control strategies for the disease (18,36). Therefore, due to the epidemiological importance of serovars related to *L. santarosai* in Colombia, we chose two highly polymorphic genes (gyrB and secY) to identify these serovars for concatenated sequence analysis. Using this methodology, it was possible to identify 26 different serovars worldwide. To date, *L. santarosai* has been associated with at least 56 different serovars (36–38). Therefore, this methodology was able to differentiate at least 46% of the currently reported serovars.

These results highlight the importance of concatenated sequence analysis to investigate the epidemiology, transmission patterns, and infection chains of L. santarosai-related serovars in Colombia. The main difficulty of this methodology was the absence of gyrB and secY gene sequences for the 56 serovars; however, this could be an attractive alternative to the complex CAAT (cross-agglutination absorption test) methodology. The usefulness of these highly polymorphic genes in differentiating species and serovars has been previously reported by Slack et al. 2006 (39) and Cerqueira et al. 2010 (40). Therefore, a combination of these genes increases the power of discrimination between serovars related to L. santarosai. The results obtained from this research are the first step to recognize the pathogenicity of the strains circulating in Colombia and facilitate the implementation of new tools for the molecular characterization of unknown Colombian isolates.

The subsequent bioinformatic analysis was the detection of orthologous proteins between the Colombian strain and the 67 species of the genus *Leptospira* together, yielding 1,650 orthologous proteins between them. This finding is interesting, because these proteins are related to the vital processes of bacteria, that is, involved in cell survival and metabolism. Another interesting finding was that most of these proteins reflect the molecular speciation process of the *Leptospira* genus. Here we describe all the orthologous genes of the Colombian strain with each of the 67 species of *Leptospira* individually, finding that the Colombian strain shares the greatest number of orthologous proteins with the subgroup of pathogenic species, *L. santarosai* being the most common species. genetically related as expected and *L. kemamanensis* the most distant species. Furthermore, with this analysis there was more similarity within the group of pathogens with the majority of pathogenic species that present greater virulence. This could coincide with the results found by Agudelo-Flórez et al. 2013 (3) which demonstrated that L. santarosai, although pathogenic, is less virulent than L. interrogans, known to have high virulence, when evaluated in an experimental model.

In the bioinformatic analysis of the detection of orthologous genes in the different groups of the genus *Leptospira* with OrthoMCL, it was found that the Colombian strain shares approximately 1,815 orthologous proteins with the representatives of the different subgroups of species, which shows that, by including more species, this central genome tends to decrease, because the greater the number, the fewer similar proteins are found among the entire genus, which helps reduce this value.

For its part, the detection of orthologous proteins between the Colombian strain and 29 serovars related to the genomes of the *L. santarosai* species available in the NCBI database determined a range of 2,820-3,491 orthologous proteins in the analyzed genomes, which shows a high genetic diversity and genomic plasticity even among *L. santarosai* isolates. This high genomic plasticity between different serovars of pathogenic species has been previously described by Ricaldi et al. 2012 (41) and Xu et al. 2016 (42), and could be related to the adaptation processes of serovars to different hosts.

On the other hand, 59.02% of the proteins were more similar to the intermediate and saprophytic species, with a percentage of identity ranging between 1 and 80%. The conservation of proteins from free-living ancestors could be related to the ability of *L. santarosai* to survive in the environment for long periods of time (5). The existence of this set of shared proteins in the *Leptospira* genus has been previously reported by Ricaldi et al. 2012 (41), and Vincent et al. 2019 (5), which supports the results found in this research.

The common and unique proteins (Table 4) in the genus *Leptospira* are ideal molecular targets for the implementation of new molecular characterization tools at the species level. Two recent studies support the existence of exclusive proteins in the genus, subgroups and different species in the genus *Leptospira*. In the first study, they performed a comparative genomic analysis to answer the question: What makes a bacterial species pathogenic? while the second study describes the pathogenesis mechanisms and virulence factors used by bacteria to cause diseases (4,43). They highlight the importance of comparative genomic analysis to describe multiple biological processes, understand genomic diversity and provide information on biological functions and evolution in the genus *Leptospira*.

The *Leptospira* genus at an evolutionary level has conserved a set of proteins among the different species and is evident in its genomic core or central genome which is conserved and closed (Fig. 6) (5,42). But at the pangenome level this is open, which also reflects the high interspecies and intraspecies genomic variability as in the case of *L. santarosai* (Fig. 10) (5,43). The open pangenome is due to the appearance of unique genes present only in some species (variable or accessory genome). This suggests great plasticity in the gene repertoire in this specific subgroup and continuous gene acquisition, which is consistent with results obtained in other studies (5,25). Contrary to closed pangenomes that remain stable and with the same number regardless of the inclusion of new species such as Bacillus anthracis and other bacteria (5,30).

Species with open pangenomes generally live in multiple environments and are in contact with mixed microbial communities, which facilitates the constant exchange of genetic material. This genomic plasticity is possibly the result of multiple ways of incorporating new genetic material into their genomes, which which increases its high variability at the genome level, allowing it to constantly expand its total gene set (5,30). This genomic diversity can be attributed to the gain by horizontal gene transfer and gene loss, reporting greater genetic losses in pathogenic strains compared to saprophytic strains, which deduces that each species makes high changes in its genetic composition to adapt to specific conditions. of variable environments. This characteristic is beneficial for the bacteria because these adaptations translate into greater infectivity in a wide range of mammals and survival in different environments (5,44).

Currently, the pathogenicity mechanisms of *Leptospira* are unknown, and this is because there are no tools for genetic manipulation studies of this bacteria (23,44). The use of genetic engineering has been the most successful methodology to evaluate bacterial virulence, and this has been carried out through the generation of random mutants and subsequent evaluation of said phenotype (44). A virulence factor is eliminated and tests are subsequently performed to quantify its level of virulence (23). This genetic silencing has been difficult to carry out in *Leptospira* because the mutations do not always occur in the gene of interest, other times these mutations fall at random in hypothetical genes of which there is little knowledge or, if they occur, they do not occur in all of them. copies of the gene of interest (44). It is unknown why it is difficult to use gene editing tools that have been successfully used in other bacteria (23). It is known that CRISPR systems have been successfully used in different bacterial groups to produce genetic editing and allow the study of virulence factors. The CRISPR-Cas silencing methodology is a novel methodology that can make up for this shortcoming and may be promising for generating mutant strains and allowing the evaluation of the different virulence factors of a certain *Leptospira* species (44).

The results obtained confirm that *L. santarosai* has a well-defined CRISPR system, as was found in other studies with other species of this genus, where said system was only located in pathogenic and intermediately pathogenic species (23,43,44). . Therefore, it is speculated that it is necessary in the production of pathogenicity, in addition to being useful to defend against exogenous factors, especially the invasion of foreign DNA or RNA (44). This finding could open several fields for pathogenicity studies in this particular species by being able to edit, through this system, specific virulence factors of this bacteria and in this way carry out delimited tests with a certain virulence factor of interest. What would be a great achievement in understanding the pathology of *Leptospira*. Although, it must be taken into account that the sequences of CRISPR systems generated by bioinformatics programs can have non-specific results, especially when short sequences that can generate matches with other CRISPR systems are included in the analysis, so experimental validation is required. to corroborate the results obtained.

Vitamin B12 (cobalamin) is the largest and most complex of the natural organometallic cofactors and coenzymes; its de novo synthesis requires about 30 enzymatic steps, which is why it is energetically expensive (4,45,46). Mammals have developed complex mechanisms to acquire, create, transport, and store this cofactor (45,46). It has been shown that only species of the *Leptospira* pathogenic group can synthesize this vitamin de novo, which allows us to speculate that it is necessary for infection processes in the host.

This bacterial genus can survive in an external environment, but only pathogenic strains can create cobalamin de novo from L-glutamate, suggesting that it is highly required in vivo. Another evidence of the above is that all pathogenic species can survive in vitro, in culture media lacking this vitamin, while saprophytic species cannot survive. One host defense mechanism is to sequester vitamin B12, similar to iron sequestration strategies (46). For this reason, pathogenic bacteria have developed these mechanisms to survive within the host, facilitating adaptation in different environments, as it allows them to live in environments lacking this vitamin (4,47).

The genes necessary for the biosynthesis of vitamin B12 in the *Leptospira* genus are grouped into two: cob I/III and cob II. It has been observed that only pathogenic leptospires are autotrophs of vitamin B12, and are capable of synthesizing it from a simple amino acid precursor such as L-glutamine (4). The necessary and precise number for such synthesis still remains unknown. Therefore, it should be noted that the exact details of this metabolic process in *Leptospira* spp are not yet known.

In the Colombian strain, an enzyme complex was found made up of 25 proteins generally involved in processes related to vitamin B12, of which 15 are directly involved with the de novo synthesis of this vitamin, which shows that it has machinery necessary to be able to infect a host, which is interesting, as it helps to understand more about this native pathogen. And it provides more support to the previous findings of this study where this bacteria has presented greater phylogenetic and evolutionary closeness to pathogenic species.

The previous comparative genomic analysis corroborated a high genomic plasticity among pathogenic species such as the Colombian strain *L. santarosai* serovar Alice, which indicates that different species manage to undergo a different evolutionary process to adapt to different hosts or environmental niches. It is important to highlight that, in a context of diagnosis and subsequent rapid classification, it is not realistic to carry out an effective and fast phylogeny based on several hundred genes, because it would be expensive, slow and wasteful. Therefore, finding an excellent candidate gene that is found in all the different groups of *Leptospira* and that allows complete differentiation of all 67 species, has to be a task on which future research efforts should focus.

For their part, the genomic characteristics evidenced bioinformatically can be basic in the knowledge of the bacterial mechanisms, the evolutionary, phylogenetic and pathogenic analysis of *Leptospira* spp, which is the basis for describing virulence factors involved in the pathophysiology of this Colombian strain and subsequently infer them in experimental models and thus delve into the knowledge of other significant aspects in the biological order of this disease that affects the population of endemic areas.

